# Structural basis for osmotic regulation of the DNA binding properties of H-NS proteins

**DOI:** 10.1101/757732

**Authors:** Liang Qin, Fredj Ben Bdira, Yann G. J. Sterckx, Alexander N. Volkov, Jocelyne Vreede, Gabriele Giachin, Peter van Schaik, Marcellus Ubbink, Remus T. Dame

## Abstract

H-NS proteins act as osmotic sensors translating changes in osmolarity into altered DNA binding properties, thus, regulating enterobacterial genome organization and genes transcription. The molecular mechanism underlying the switching process and its conservation among H-NS family members remains elusive.

Here, we focus on the H-NS family protein MvaT from *P. aeruginosa* and demonstrate experimentally that its protomer exists in two different conformations, corresponding to two different functional states. In the half-opened state (dominant at low salt) the protein forms filaments along DNA, in the fully opened state (dominant at high salt) the protein bridges DNA. This switching is a direct effect of ionic strengths on electrostatic interactions between the appositively charged DNA binding and N-terminal domains of MvaT. The asymmetric charge distribution and intramolecular interactions are conserved among the H-NS family of proteins. Therefore, our study establishes a general paradigm for the molecular mechanistic basis of the osmosensitivity of H-NS proteins.

## Introduction

Bacteria are frequently exposed to changes in their extracellular *milieu*. Key to their survival is to adapt quickly to these environmental fluctuations by adjusting their cellular machinery, enabling them to operate optimally under the new conditions, and to acquire new competencies through horizontal gene transfer (HGT) ^1^. These adaptation mechanisms are regulated by the bacterial genome, the dynamic organisation of which modulates access to the genetic material to adapt and fine-tune gene expression and for incorporation of DNA of foreign origin ^2^.

The Histone-like nucleoid structuring (H-NS) protein is a key regulator of the dynamic bacterial genome ^3, 4^. The protein is conserved among enterobacteria and plays determinant role in their nucleoid architecture. Generally, this protein acts as repressor of transcription, silencing the expression of many different genes and operons. Often the genes and operons targeted by H-NS family proteins are regulated by environmental factors such as osmolarity, pH and temperature ^5, 6^. Additionally, H-NS family members have the unique ability to “switch off” xenogeneic genes, often in pathogens, to suppress their detrimental effect on bacterial fitness, until external conditions require expression ^7, 8, 9, 10^. To similar effect, H-NS has been shown to silence spurious transcription at intragenic promoters in *E. coli* ^11^.

H-NS is expressed at high levels (> 20 000 copies per cell). The protein binds non-specifically to DNA, but has a preference towards genomic regions with high AT content ^9, 10, 12^, containing specific six contiguous steps^13^. DNA binding typically initiates at such high-affinity sites on the DNA ^14^. This step is followed by oligomerization of the protein, forming lateral nucleoprotein filaments ^15^, which stiffens DNA ^16^. Under appropriate conditions the protein can mediate bridge formation between this filament and a second DNA duplex ^17, 18, 19, 20, 21^.

The current paradigm is that divalent ions drive the transition between lateral H-NS-DNA filaments and bridged DNA-H-NS-DNA complexes, without dissociation of the protein from the DNA ^19, 22^. This switching between the two types of complexes is believed to be the mechanistic basis of the role of H-NS proteins in bacterial nucleoid organisation and transcription regulation ^19, 22, 23^. Nonetheless, the molecular basis that governs this phenomenon is still controversial. Previously published molecular dynamics of H-NS, indicated that the switch between the two DNA binding modes involves a change from a half-open (also referred as a closed conformation) to an open conformation driven by MgCl_2_ ^19^. These conformational changes are modulated by the interactions between the N-terminal domain of H-NS and its C-terminal DNA binding domain (DBD). Mutagenesis at the interface of these domains generated an H-NS variant no longer sensitive to MgCl_2_, which can form filaments and bridge DNA. Recently, these interactions were confirmed by Arold and co-workers using H-NS truncated domains ^24^. However, it was claimed that these interactions could not occur unless H-NS oligomerization is abolished by temperature. These discrepancies in conclusions arise presumably from the lack of experimental structural information on the full-length protein in solution.

H-NS family members share a conserved fold topology, with a C-terminal DBD and an N-terminal domain connected via a flexible linker ^25, 26^. The structural unit of an H-NS nucleoprotein filament is a dimer (also called protomer) formed through the N-terminal dimerization sites of two monomers (site 1), which further oligomerize through a secondary oligomerization site (site 2) ^27^ (Fig. 1 a,b). Multiple studies have revealed the structures of H-NS truncated domains including the dimerization sites and the DBD from different bacteria ^13, 28, 29, 30^. Meanwhile, no structure of the full-length protein is available. The structure of the N-terminal domain oligomer (1-83) of the *S. typhimurium* H-NS C21S mutant was solved by crystallography^31^. These studies revealed a helical scaffold of a head-head and tail-tail protomer organisation, suggesting structural models involving that type of H-NS arrangement in both lateral H-NS-DNA filaments and bridged DNA-H-NS-DNA complexes.

**Figure 1:**
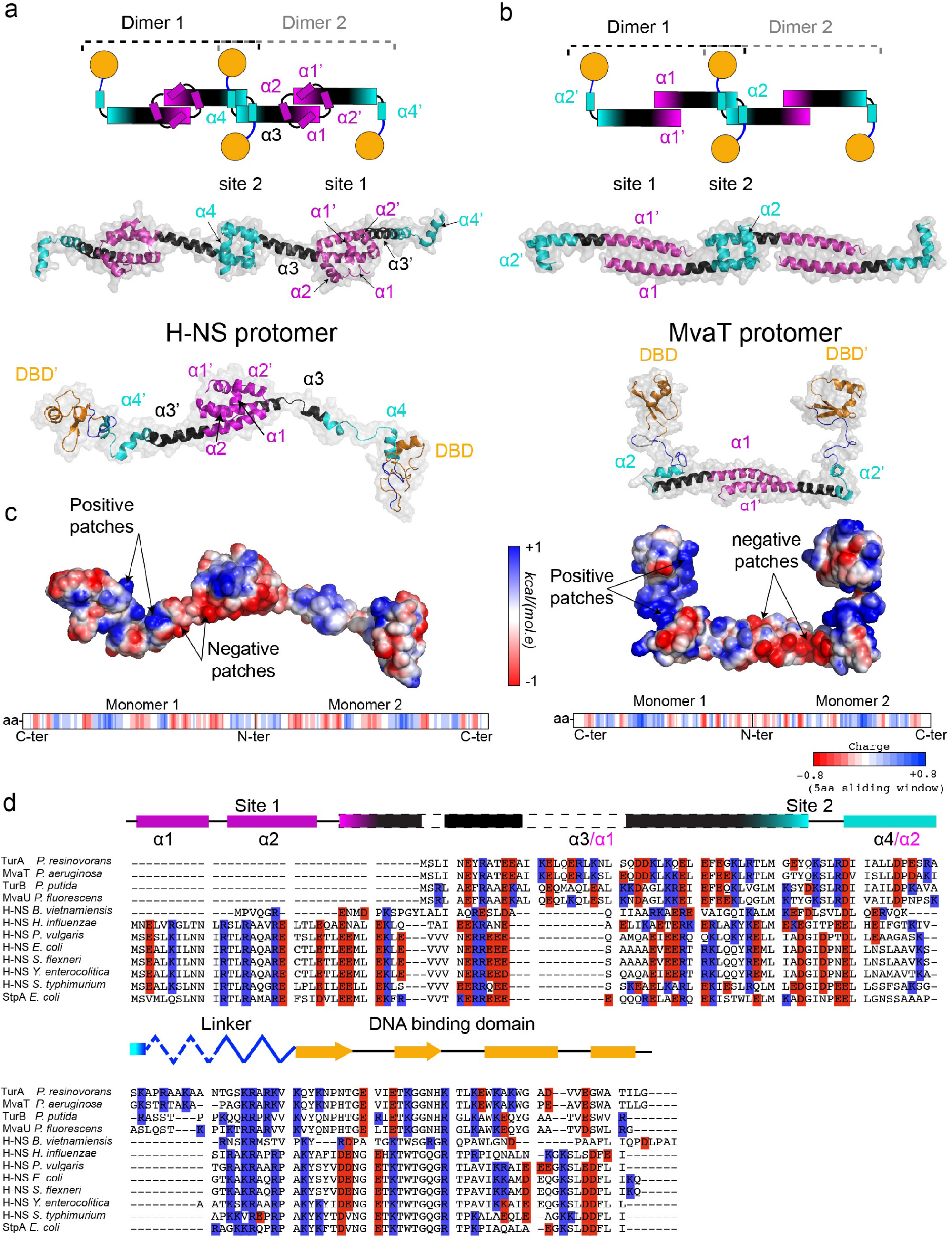
Fold topology and charged residue/charge distributions of H-NS family proteins. (a) N-terminal dimerization sites of long H-NS family members. The upper panel depicts a schematic representation of the long H-NS oligomer. The dimerization sites (site 1) are shown in magenta and the oligomerization sites (site 2) in cyan. The DNA binding domain is shown in orange sphere and the linker region in blue. In the middle panel is the H-NS N-terminal domain crystal structure (1-83) (PDB ID: 3NR7) ^31^. Site 1 and Site 2 helices are coloured and labelled as in the upper panel. In the lower panel is the H-NS protomer structural model adapted from our earlier MD simulation studies ^19^. (b) N-terminal dimerization sites of short H-NS family members. The upper panel depicts a schematic representation of the short H-NS oligomer. In the middle panel is the TurB N-terminal domain crystal structure (1-64) (PDB ID: 5B52) ^39^. Lower panel is the MvaT protomer structural homology model (see M&M). The colour code is as in (a). (c) The electrostatic potential surfaces of H-NS and MvaT are depicted on full-length protomer structural models of the proteins using a red/white/blue colour gradient. The negative and positive patches on the proteins’ surfaces are indicated. In the lower panel is the five amino acid averaged charge window of the proteins protomers primary sequences. Positive, negative and neutral charged amino acid fragments are shown in blue, red and white bars, respectively. (d) Primary sequence alignment of selected members of the H-NS family of proteins from different bacterial organisms. The conserved positions of the charged amino acids in the protein’s primary sequences are highlighted in blue (positive residues) in red (negative residues). The fold diagram of the H-NS monomer from *E. coli* is represented using the same colour code in (a). The nomenclature of the N-terminal helices is indicated in black for the long H-NS family members and in magenta for the short members.

Early studies by liquid state NMR on full length H-NS from *E. coli* and *S. typhimurium* have shown that resonances of the N-terminal domain residues were missing in the protein heteronuclear single quantum coherence (HSQC) NMR spectrum ^28, 32^. A similar observation was made using solid-state NMR in which case only 33% of the protein spectrum was assigned ^33^. These difficulties were attributed to the inherent dynamics and structural heterogeneity of the system. Thus, owing to their flexibility and the large size of their self-assembled complexes, the characterization of the conformational landscape of full-length H-NS proteins and its contribution to functional regulation remains a challenge. In this work we have overcome these constraints by using an abridged structural version of H-NS, MvaT from *P. aeruginosa*, in which we scrutinized its protomer structural/function relationship in response to changes in the surrounding ionic strength.

MvaT is a functional homologue of H-NS that regulates the expression of ~350 genes including the genes responsible for virulence ^34^ and biofilm formation ^35, 36^. Like other members of the H-NS family, oligomerization of MvaT along DNA is required for its gene silencing activity ^37, 38^ and ability to stiffen and bridge DNA duplexes ^18, 38^.

Here, we have combined integrative structural biology methods and (single-molecule) biochemical assays to decipher the structural changes in MvaT that drive the switch between its DNA stiffening and bridging activities under the influence of different salt conditions. These structural changes appear to be conserved within the H-NS family of proteins. Our studies thus provide a fundamental answer on how H-NS family members respond to changes in the osmotic environment to modulate their DNA binding modes, and hence to regulate bacterial nucleoid organisation and gene expression.

## Results

### Fold topology and electrostatics of H-NS family proteins

Primary sequence alignments and secondary structure predictions of twelve members of the H-NS family of proteins, from different bacteria, reveal a conserved fold topology (fig. S1). Their C-terminal DBD contains, in most cases, two β-strands and two α-helices. The domain is connected to an N-terminal domain via a disordered linker. The N-terminal domain contains α-helices connected via random coils, but the number of helices may differ. Based on these criteria, two classes of H-NS members could be distinguished: long and short versions of H-NS. The long H-NS members have an N-terminal domain which includes four helices (α1, α2, α3 and α4), forming a dimerization site (site 1) between the H-NS monomers and an oligomerization site (site 2) between its protomers (fig. 1a). The crystal structure of the H-NS N-terminal domain (1-83) has shown that site 1 is formed by a “hand-shake fold” topology between α1 and α2 and part of α3. Site 1 is stabilized by a hydrophobic core and salt bridges ^30, 31^ (fig. 1a). In the short members (MvaT; MvaU; TurA and TurB), these two N-terminal helices are missing (fig. 1b; S1). The crystal structure of the TurB N-terminal domain (1- 61) has revealed that site 1 is formed by a “coiled-coil motif” between the N-terminal helices α1 (corresponding to H-NS α3) of two monomers ^39^ (fig. 1b).

Using the TurB N-terminal domain crystal structure, we have built a homology model of the MvaT dimerization site which was connected via the flexible linkers to the NMR structure of the protein DNA binding domains (PDB ID: 2MXE) ^40^, thus obtaining a full-length structural model of the MvaT protomer (fig. 1b). We used this model for structural investigation.

In contrast to the differences in site 1, the oligomerisation site (site 2) fold architecture appears to be conserved between these members of the H-NS family (fig. 1a,b). This site is also formed by hydrophobic interactions between two α-helices (α4 and α2 for the long and short members, respectively) and stabilised by salt bridges ^31, 39^.

One striking characteristic is a local degree of conservation of the electrostatic potential between H-NS and MvaT protomers. Their electrostatic potential surfaces show that the N-terminal domain (helices α3 and α1 of H-NS and MvaT, respectively) contain negatively charged surface patches, whereas the C-terminal domain and the linker region contain positively charged regions (fig. 1c).

This characteristic appears to be shared between the H-NS family members, as it can be predicted from the conserved position of the negatively and positively charged residues within their primary sequences (fig. 1d). The five amino acid averaged charge window analysis of the primary sequences revealed that the α-helices in the N-terminal domain include predominantly negatively charged regions, while the linkers and the DNA binding domains are mostly positively charged, as they both contribute to DNA binding ^41, 42, 43^ (fig. 1d; S1). Additionally, α1 and α2 of the long H-NS members are positively charged in line with their contribution to the interaction with DNA ^30^ (fig. 1d; S1). Thus, this asymmetrical charge distribution seems to be a conserved feature between H-NS family members suggesting a specific role in their function.

In solution, it is expected that the two DNA binding domains of the H-NS protomers behave as ‘beads on a flexible string’, where the function of the linkers is to enable a relatively unhindered spatial search by the attached domains. Thus, electrostatic interactions between the oppositely charged domains, as suggested by earlier studies^19, 24^, might occur. These interactions are possibly sensitive to changes of ionic strengths in the protein environment.

### Salt induces global conformational changes in MvaT protomer

To test our hypothesis that the MvaT protomer conformation is sensitive to changes in ionic strength of the surrounding medium, we employed a combination of circular dichroism, small-angle X-ray scattering (SAXS) and NMR spectroscopy. To this purpose, we engineered an MvaT variant in which oligomerization was abolished by substituting site 2 residues F36 and M44 with aspartic acid (D), remote from the negative charge patch of the N-terminal region. As site 1 is left unchanged, it was expected that MvaT could still form dimers, which was confirmed by size-exclusion chromatography (fig. S2a,b). The double mutation induces changes in the MvaT α-helical content as concluded from the overlay between the wild type and the mutant far UV CD spectra (fig. S2c). A similar observation was reported for H-NS from *S. typhimurium*, which was attributed to the loss of the protein oligomerization ability^32^. The effect of the double mutation on the binding of MvaT to DNA was assessed by electrophoretic mobility shift assay (fig. S2d,e). Binding of the MvaT wild type caused a major reduction in DNA substrate mobility, whereas for the double mutant DNA mobility was only mildly affected. We attribute this to a lack of DNA binding cooperativity for the dimeric MvaT derivative, as confirmed by tethered particle motion (see M&M) (fig. S2f).

SAXS was employed to assess the effect of salt concentration on the global conformation of the MvaT F36D/M44D dimer (fig. 2a and Table S1). Under low salt conditions (50 mM KCl) the MvaT protomer adopts a relatively compact conformation as evidenced by the normalised Kratky plot and the particle’s Porod exponent (3.2) (Table S1). Interestingly, at high salt concentrations (300 mM KCl), the MvaT protomer displays an increased flexibility, which is reflected by an increase in R_g_ (3.56 nm vs. 3.83 nm) and D_max_ (14.70 nm vs. 15.84 nm), a decrease in Porod exponent (3.2 vs. 2.7), and the appearance of the normalised Kratky plot (fig. 2a and Table S1).

**Figure 2:**
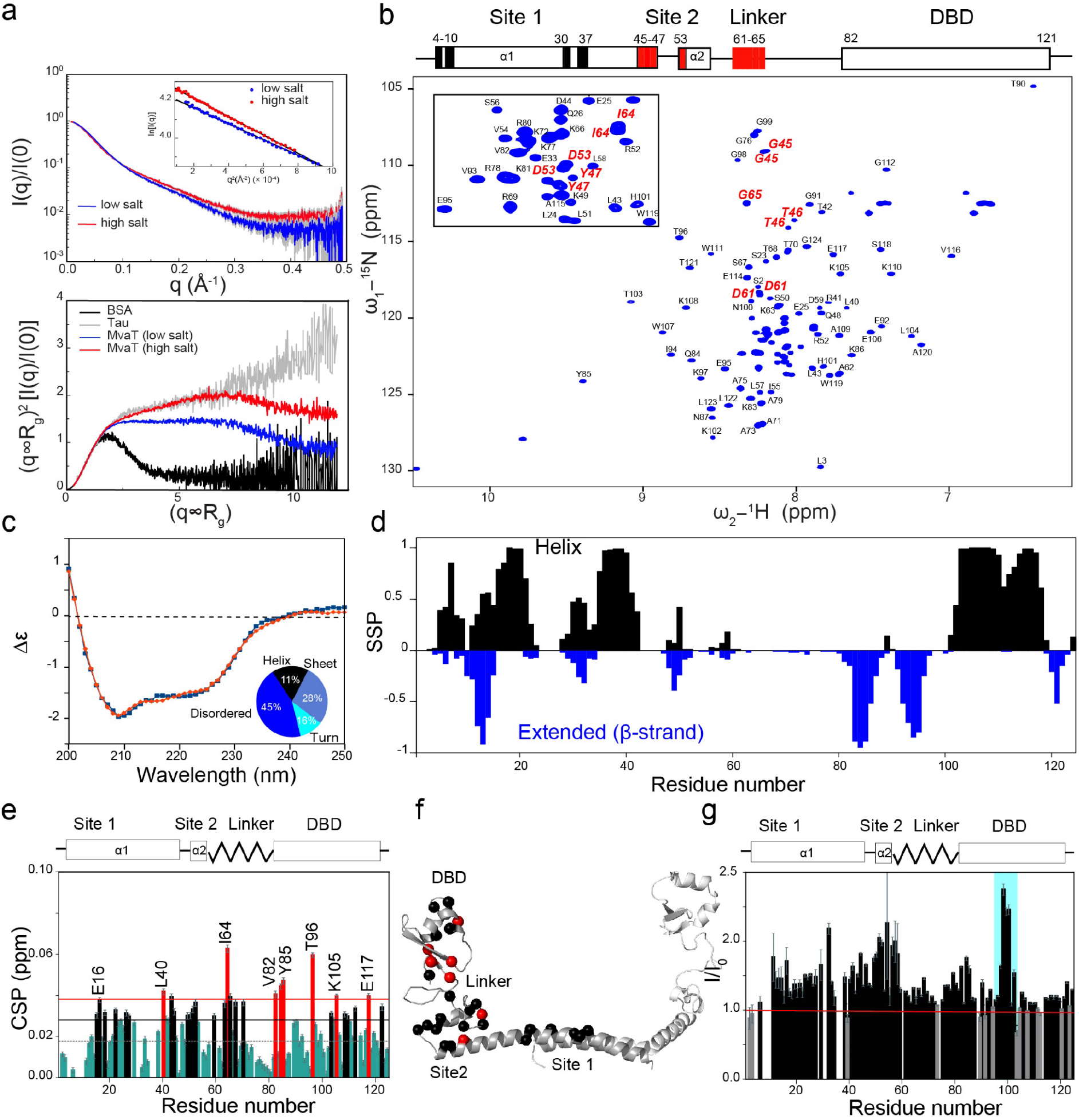
Global conformational changes in MvaT induced by salt. (a) SAXS experiments. The top panel displays the SAXS data obtained for the MvaT dimer under low salt (50 mM KCl) and high salt (300 mM KCl) conditions indicated by the blue and red curves, respectively. The error margins are coloured in grey. The inset shows the Guinier regions of both data sets. The bottom panel displays the normalised Kratky plots for the MvaT dimer under both salt conditions. An intrinsically disordered standard (Tau, grey trace) and a rigid, globular standard (BSA, black trace) are shown for comparison. (b) Assignment of the ^1^H-^15^N HSQC MvaT spectrum (at 150 mM KCl). Residues that appear as double peaks in the spectrum are labelled in red. Insert is a zoom in at the crowded region of the spectrum. The layout of the MvaT domains is shown above the spectrum. Residue positions with non-observable resonances in the HSQC spectrum are marked by black rectangles. The residue positions with two observable resonances are in red. (c) Circular dichroism spectra of MvaT dimer (experimental data in blue and spectrum deconvoluted using BeStSel in red ^44^). (d) Secondary structure probability (SSP) plot of the MvaT dimer derived from Cα and Cβ chemical shifts, determined by PECAN^45^. Negative values in blue bars indicate β-strand or extended secondary structures and positive values in black bars indicate α-helices. (e) Weighted average CSP of the MvaT resonances between 50 and 300 mM KCl. Resonances with CSP more than two (red line) or one (black line) standard deviation (SD) from the 10% trimmed mean (grey dashed line) are labelled and shown in red, black and green bars, respectively. (f) Residues exhibiting CSP mapped on the structural model of the MvaT dimer (see M&M). Amide nitrogens with a CSP > 1 and 2 SD trimmed mean are shown in spheres and coloured in black and red, respectively. (g) The HSQC peak intensity ratio of the MvaT dimer at 300 mM KCl (I) and 50 mM KCl (I_0_). Grey bars indicate residues with minor peak intensity changes between the two salt conditions. Loop 95-102 of the DBD is highlighted with a cyan rectangle.

In parallel to the SAXS analysis, we produced an isotopically labelled ^15^N ^13^C MvaT sample for which the ^1^H-^15^N HSQC NMR spectrum was assigned at 150 mM KCl. The protein spectrum shows well-dispersed resonances corresponding to residues of the DBD and multiple overlapped peaks (crowded region) centred around 8.2 ppm in the ^1^H dimension, reporting on the helical N-terminal domain and the disordered linker (fig. 2b). Remarkably, the MvaT spectrum shows more resonances than expected, indicating the presence of two forms. Two resonances are observed for the amide groups of residues 45-47 and residue 53, which form the oligomerization site (site 2), and of residues 61-65 in the linker region (fig. 2b). Amides of the “coiled-coil” (site 1) show weak resonances and several of them could not be assigned due to broad lines (fig. 2b). The Cα and Cβ chemical shifts were used to predict the secondary structure probability of the MvaT dimer in solution (fig. 2d). Amino acid sequences of α-helix 2 (51-58), the linker (60-80) and of short stretches of α1 appear to be disordered. The presence of intrinsically disordered regions agrees with the secondary structure prediction from the far-UV CD spectrum. The deconvolution of the protein CD spectrum revealed that 40-45 % of the protein is disordered (fig. 2c).

Next, we investigated the binding of potassium chloride to the MvaT protomer by NMR titration. Small chemical shift perturbations (CSP) in the fast exchange regime were observed for multiple resonances. Residues exhibiting CSP are scattered on regions of site 1, site 2, the linker and the DBD in the structural model of the MvaT dimer (fig. 2f), suggesting a non-specific effect due to the increase in ionic strength. Interestingly, the protein resonances show a general yet non-uniform increase of intensities at high salt compared to low salt. This suggests a decrease in the rotational correlation time of the protein folded domains (fig. 2g), possibly caused by the disruption of their intramolecular interactions. This observation is in line with the larger radius of gyration and the enhancement of protein flexibility derived from the SAXS data.

In conclusion, the increase in ionic strength, indeed induces global conformational changes of the MvaT dimer by shifting its conformational equilibrium from relatively compact to extended and flexible states. These structural changes are possibly due to the salt modulation of the postulated intramolecular electrostatic interactions between the oppositely charged regions of the MvaT protomer.

### Salt modulation of the MvaT dimer conformational landscape

To obtain more information about the conformational changes induced by salt on the MvaT dimer, we used paramagnetic relaxation enhancement (PRE) NMR spectroscopy. PRE reports on long-range structural distances (20 to 30 Å) between a paramagnetic spin label and the amide protons of the protein ^46^. Amide protons on residues within 30 Å from the spin label experience an increase in relaxation rate resulting in broadening of their NMR resonances and intensity reduction. To measure PRE on the MvaT dimer, we substituted residue K31 with a cysteine and conjugated it with either a paramagnetic (MTSL) or a diamagnetic (MTS) probe (see M&M).

Under low salt condition (50 mM KCl), strong PRE effects were observed for residues of site 1 (1-48) (fig. 3a). These residues were excluded from the analysis as they display weak peak intensities in the HSQC diamagnetic spectrum and sense the paramagnetic effects from spin labels on both MvaT subunits. Residues of site 2 (49-60) and the linker region (61-80) exhibit also strong PREs, indicating that the amide nuclei spend at least part of the time close the probe (fig. 3a). A weak PRE effect occurred for residues of the DBD, except for the loop between 95-105 that, in the DNA-bound state, is known to intercalate with the DNA minor groove ^40^. The strong PRE effects within this loop suggest a rather localized interaction between the DBD and a region of the protein close to the spin label.

**Figure 3:**
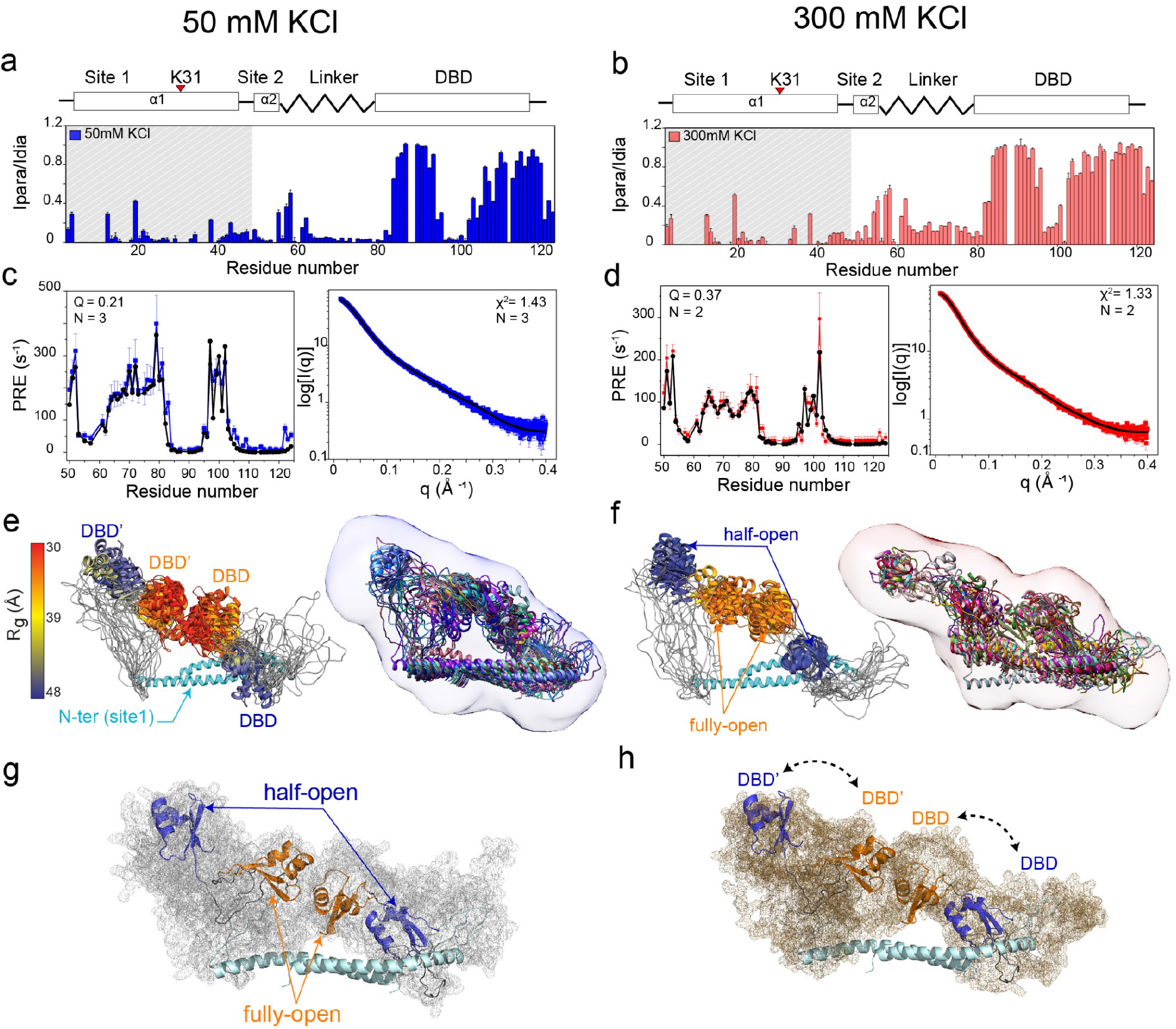
MvaT dimer structural ensembles at low and high salt conditions. (a), (b) The PRE-intensity ratio (Ipara/Idia) vs. residues number of the MvaT dimer at 50 mM KCl (blue bars) and 300 mM KCl (red bars), respectively. PRE from residues in the shaded region were excluded from the analysis. The structure of the MvaT monomer is shown schematically above the panels and the MTSL labelling position is indicated by the red triangle. The error bars are inversely proportional to the propagated signal-to-noise ratio of individual resonances. (c) and (d) The experimental PRE and SAXS curves (50 mM KCl in blue and 300 mM KCl in red) fit (black line) to the top conformers of the low and high salt ensembles, respectively. (e, f) Low (e) and high salt (f) best conformers coloured by their radius of gyration using a blue/orange gradient, respectively. The colour gradient is depicted only on the DBD of the MvaT dimer (DBD’ and DBD refer to MvaT subunits 1 and 2, respectively). The best conformers are fitted into the *ab initio* molecular shapes generated by DAMMIF ^47^. The low (g) and high salt (h) half-open and fully open conformers of the MvaT dimer, classified based on the position of DBDs relative to the N-terminal domain. The atomic probability distribution maps of the low and high salt ensembles are shown in grey and brown meshes, respectively. The DBDs are shown in blue for the half-open conformers and in orange for the fully-open conformers. The dashed arrows indicate the exchange between the DBD configurations.

At 300 mM KCl, the PRE pattern is overall similar, but a decrease in PRE is observed for the site 2 residues, the linker regions and loop 95-105 of the DBD (fig. 3b). Under this condition, these regions could either be populating conformations in which the nuclei are further away from the spin label or spend a smaller fraction of the time in the conformation close to the spin label.

To investigate the structural states sampled by the MvaT dimer under both salt conditions, we performed ensemble calculation using an Xplor-NIH protocol which combines the PRE and SAXS data as structural restraints. For each condition, we simulated molecular ensembles of multiple conformers (N = 1-5) and assessed their agreement with the experimental data by calculating the corresponding quality metrics (PRE Q-factor and the SAXS χ^2^) (see M&M). The best solutions for the low salt were obtained with N=3 (Q = 0.21; χ^2^ = 1.43) (fig. 3c; S3a) and for the high salt with N=2 (Q = 0.37; χ^2^ = 1.33) (fig. 3d; S3b). The 10 lowest-energy solutions for each ensemble fit well within the molecular shapes determined by DAMMIF (fig. 3e,f) ^47^.

In agreement with the flexibility observed for MvaT in both low and high salt conditions (with a significantly more pronounced flexibility at high salt concentration), the SAXS and PRE data sets could only be described by a conformational ensemble. As expected, under low-salt conditions the MvaT ensemble is composed of rather compact conformers, while at higher salt concentrations the ensemble members adopt more extended conformations due to an increased flexibility (fig. 3e,f).

Based on the DBD position relative to the N-terminal domain two major species could be distinguished in both conformational ensembles: half-opened and fully-opened forms. In the half-opened states, the DBD is bound to the N-terminal domain, while the other domain is free (fig. 3g,h). The occupancy of the N-terminal domain with one of the DBD of the dimer appears to prevent simultaneous binding of the second DBD to the N-terminal domain due to steric hinderance. This might explain the observed asymmetrical structural configuration of the MvaT homodimer in the half-open forms.

In the fully-opened states, both DBDs are not bound to the N-terminal domain but can still be close to each other in space. Both of conformational species comprise several sub-populations with different radii of gyration (fig. 3e,f).

Under both salt conditions, the DBD in the major conformer of the half-opened states exhibits similar intra-subunit binding modes to the N-terminal domain, although a slight difference in orientation is observed (fig. S3c). The binding interface includes the positively charged loop 95-ET**K**GGN**HK**-102 (charge = 1.8, pH 6) of the DBD and the negatively charged segment 26-Q**DD**KLKK**E**L**EDEE**-38 (charge = −3.8, pH 6) of the N-terminal domain. In contrast, at low salt the linker appears to form closer interactions with the negatively charged patch of the N-terminal domain (26-38) compared to the high salt condition. This indicates that the linker might be involved in stabilizing the DBD-N-terminal domain complex.

In conclusion, the structural study reveals the existence of electrostatic interdomain interactions between the DBD and N-terminal domains of MvaT which are further stabilized by the linker region. Increase in salt concentration induces delicate changes within the MvaT protomer conformational landscape by weakening these intramolecular interactions and increasing the exchange rate between the half-open and open conformers. Consequently, the protein samples a larger conformational space.

### The DNA substrate contributes to the MvaT protomer conformational opening

Owing to the negative charge and specificity for the DBD it is conceivable that the DNA also modulates the interdomain interactions of the MvaT protomer. To test this hypothesis, a titration of the MvaT dimer with a 12 bp AT-rich DNA substrate at low (50 mM KCl) and high salt (300 mM KCl) concentrations, was performed using NMR. At low salt, significant CSP were observed for residues of the DBD. At the DNA:MvaT molar ratio of 1.6, a general line broadening of the DBD peaks is observed attributed to the increase of its rotational correlation time (τ_c_) upon complex formation, independent of the protomer N-terminal domain (fig. S4a). Analysis of the titration data using a 1:1 binding model yields a K_D_ = 44 ± 5 nM and k_off_ = 280 ± 5 s^−1^ (fig. S4c). The affinity is too high to be deduced from this NMR titration with accuracy. Therefore, we used ITC for validation, yielding K_D_ = 30 ± 2 nM (fig. S4d). No significant CSP for the N-terminal domain residues were observed, indicating that the interactions with the short DNA substrate are limited to the DBD and a few residues of the linker region (fig. 4a). However, a 1:1 binding model may be an oversimplification.

**Figure 4:**
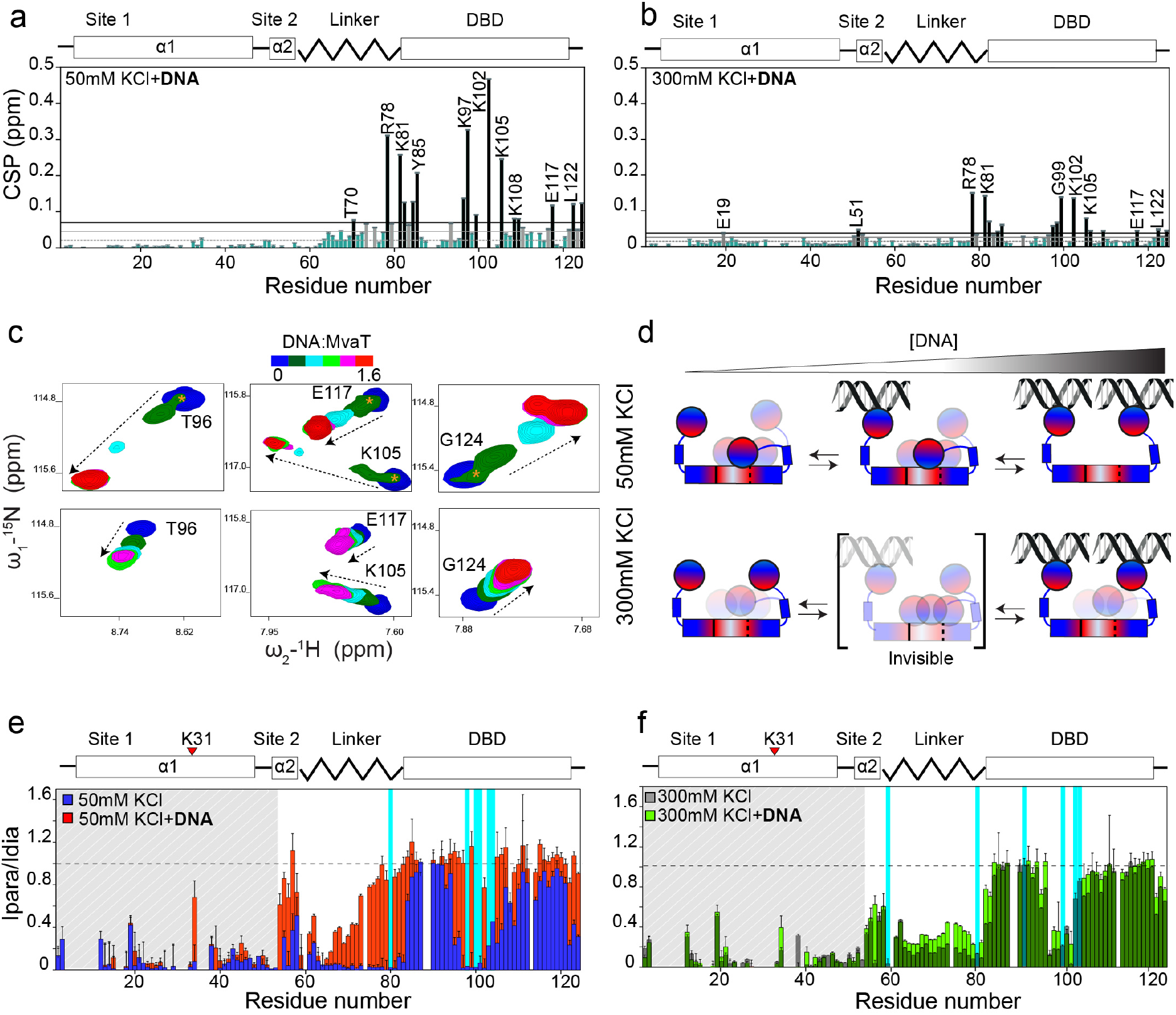
DNA modulates the conformational state of the MvaT dimer. (a) and (b) Analysis of the MvaT dimer HSQC spectra weighted average CSP at 1.6 DNA:MvaT molar ratio at 50 and 300 mM KCl, respectively. Resonances with CSP more than two (black line) or one (grey line) standard deviation (SD) from the 10% trimmed mean (grey dashed line) are labelled and shown in black, grey and green bars, respectively. (c) CSP amplitudes and directions (dashed arrows) of the DBD resonances upon titration with 12 bp DNA at 50 (upper panels) and 300 mM KCl (lower panels). The resonances that correspond to the N-terminal bound DBD are indicated by orange asterisks. (d) Schematic representation of the DNA binding mechanism to the MvaT dimer under low and high salt conditions. The electrostatic characteristics of the MvaT dimer are schematically shown by a red /white /blue colour gradient. (e) and (f) Overlay between the PRE vs. MvaT dimer residue number at low salt in the absence (blue bars) and presence of DNA (red bars) and high salt in absence (grey bars) and presence of DNA (green bars), respectively. Resonances with non-determined PRE, due to line broadening upon DNA binding, are highlighted in cyan bars. The layout of the MvaT monomer is shown in the upper panel. The position of the spin label is indicated by the red triangle.

The ensemble structure of the MvaT dimer at 50 mM KCl revealed that interactions between the DBD and the N-terminal domain occur, with only one of the two DBD binding at any time, effectively creating two populations of DBD (fig. 4d). Thus, one DBD is free to bind to the DNA, whereas for the other competition for binding occurs between DNA and the N-terminal domain. The NMR spectra seem to catch this difference in the DBD forms at a DNA:MvaT molar ratio of 0.2, showing two peaks with different CSP for the DBD resonances (fig. 3e; 4c). In the absence of DNA, the DBD will be in exchange between bound and free states, and also within the bound state, it may sample multiple interactions in an encounter complex like state. Broad lines are observed for multiple amide resonances including the ones that correspond to residues of loop 95-105 (fig. S3d; 1g), suggesting that these exchange processes are intermediate to fast on the NMR time scale (i.e. the chemical shift differences between bound and free). Addition of DNA will bind the population of free DBD preferably, splitting the resonances of the two populations, provided the exchange rate between the states of DBD bound to the N-terminal domain and bound to the DNA is sufficiently slow. At higher concentrations of DNA the exchange rate is increased (k_ex_ = k_on_[DNA] + k_off_) and the exchange between the two DBD populations is too fast to result in double peaks. This model is supported by titration studies of the isolated DBD with DNA. In the absence of the N-terminal domain, the resonances of the DBD show single peaks at 0.1 and 0.2 DNA:MvaT molar ratios (fig. S3e).

At high salt, the interdomain interactions are weakened by the ionic strength and the exchange rates between the DBD free and bound to the N-terminal becomes fast on the NMR time scale. Therefore, the peak splitting, observed under the low salt condition, becomes invisible (fig. 4c,d). Interestingly, in this case smaller amplitude of CSP are observed upon DNA binding, yet with the same direction as for the low salt (fig. 4b). This suggests that part of the complex is in an encounter state which is in equilibrium with the specific state and does not contribute to CSP^48^. Nonetheless, a general line broadening of the DBD resonances remained observable (fig. S4b) and residues involved in the direct interaction with the DNA disappear. Analysis of the titration data yields a K_D_ = 7.7 ± 0.3 μM and a k_off_ = 2 ± 0.2 × 10^3^ s^−1^ (fig. S4b,c).

The conformational opening of the MvaT protomer induced by DNA, at low and high salt, was probed by changes in the PRE upon addition of DNA (at DNA:MvaT = 1.6). At low salt, a substantial loss in the PRE is observed for residues 60-80 of the linker region and the dimerization site 2 (fig. 4e). Almost no PRE effect on the DBD was observed. This indicates that the protein is adopting a fully-opened conformation in which the DNA substrate is shielding the DBD and part of the linker region from the interaction with the N-terminal domain. At high salt, this effect is less pronounced, yet DNA still reduces the PRE (fig. 4f). These results provide evidence that salt and DNA substrate have an additive effect on the conformational opening of the MvaT protomer.

### Modulation of MvaT electrostatic interdomain interactions by salt and DNA drives the switch between its DNA binding modes

Our structural studies have established that the MvaT dimer samples a heterogenous conformational space dominated by half open and fully open conformers. Increase in salt and DNA concentrations destabilizes the electrostatic interactions between the DBD and the N-terminal domain. Consequently, the exchange rate between the half open and fully open conformers becomes fast and the sequestration of the DBD by the N-terminal domain is diminished. Thus, the protomers can potentially interact simultaneously with two independent DNA segments.

To test whether this dynamic conformational equilibrium has implications for the structure and function of MvaT wild type-DNA complexes, we determined the modes of DNA binding under the low and high salt conditions using tethered particle motion (TPM) and bridging pull-down assay ^19, 49^.

First, we investigated the effect of salt on lateral nucleoprotein formation by oligomerization of MvaT along DNA. In TPM the length and stiffness of the tethered DNA molecules is correlated to the root mean square displacement (RMS) of the attached polystyrene bead. The formation of a filament along DNA results in DNA stiffening, observed as an increase in RMS by TPM. The RMS changes of the polystyrene bead attached to a single DNA molecule (685 bp) were determined as a function of protein concentration. The experiment was performed at 50 mM KCl (low salt) and 315 mM KCl (high salt) (fig. 6a). Under both conditions two binding regimes were evident. At low MvaT concentration, a decrease in RMS was observed, which is attributed to the ability of MvaT to bend double stranded DNA (phase I) ^40^. We propose that this regime represents a nucleation phase where MvaT protomers bind non-cooperatively to the DNA substrate (fig. 6c). In the second regime (II) a gradual increase in RMS occurs, which reaches a maximum of 160 nm at 1.6 μM of MvaT. This represents a phase in which the MvaT protomers gradually oligomerize along the DNA until saturation. The difference between the two salt conditions is in the first regime in which the DNA is bent by individual dimers (I). At high salt, the extent of DNA compaction resulting from DNA bending by MvaT is reduced (fig. 6a). This could be due to an effect of the counterions (Cl^−^) on the DNA binding affinity of MvaT (as seen with H-NS)^19^. To minimize this potential counterion effect, we also performed the TPM experiment using potassium glutamate (Kglu) instead of KCl. Indeed, the differences in the MvaT titration curves obtained at low and high salt were much less pronounced than with KCl (fig. 6b).

These results suggest that the low and high salt conditions have no effect on the MvaT DNA stiffening activity. At high salt, the delicate conformational changes due to the increase in flexibility and in exchange rates between the a half-open to a fully open states of MvaT protomers have no influence on this process. Besides, the dissociation effect of the increase in ionic strength on the complex is countered by the cooperative binding of the MvaT protomers along the DNA.

Previously we have demonstrated that H-NS switches between its DNA stiffening and bridging activities as function of MgCl_2_ by using a sensitive and quantitative bridging pull-down assay^47^. Here we have used the same approach to test the effect of potassium chloride concentrations on MvaT-mediated DNA-DNA bridging. The DNA bridging efficiency (i.e. the percentage recovery of DNA from solution) as a function of KCl concentration (fig. 6d) revealed that at low salt concentration (50 mM KCl) the MvaT-mediated bridging activity is negligible despite the protein’s ability to stiffen DNA by lateral nucleoprotein filament formation (fig. 6a). This agrees with our structural model in which MvaT protomers can bind to the DNA with only one of their DBDs while the other one is sequestered by the electrostatic interaction with the N-terminal domain. Under this condition the lateral nucleoprotein filaments of MvaT cannot form a bridge between two DNA duplexes.

By increasing the salt concentration, an increase of the protein bridging efficiency is observed and reaches a maximum at 315 mM KCl (Phase I’). This dependency is also observed in the presence of Kglu and NaCl (fig. 6d, f). The low and high salt conditions have no effect on the protein oligomerization state (fig. S5a).

We have also tested the effect of divalent ions on the MvaT DNA bridging activity using MgCl_2_, MgSO_4_ and CaCl_2_: the DNA bridging efficiency of MvaT also depends on the tested divalent ions. However, optimum bridging efficiency is reached at lower salt concentrations (~18 mM for MgCl_2_ and ~20 mM for MgSO_4_ and CaCl_2_) (fig. 5e).

**Figure 5:**
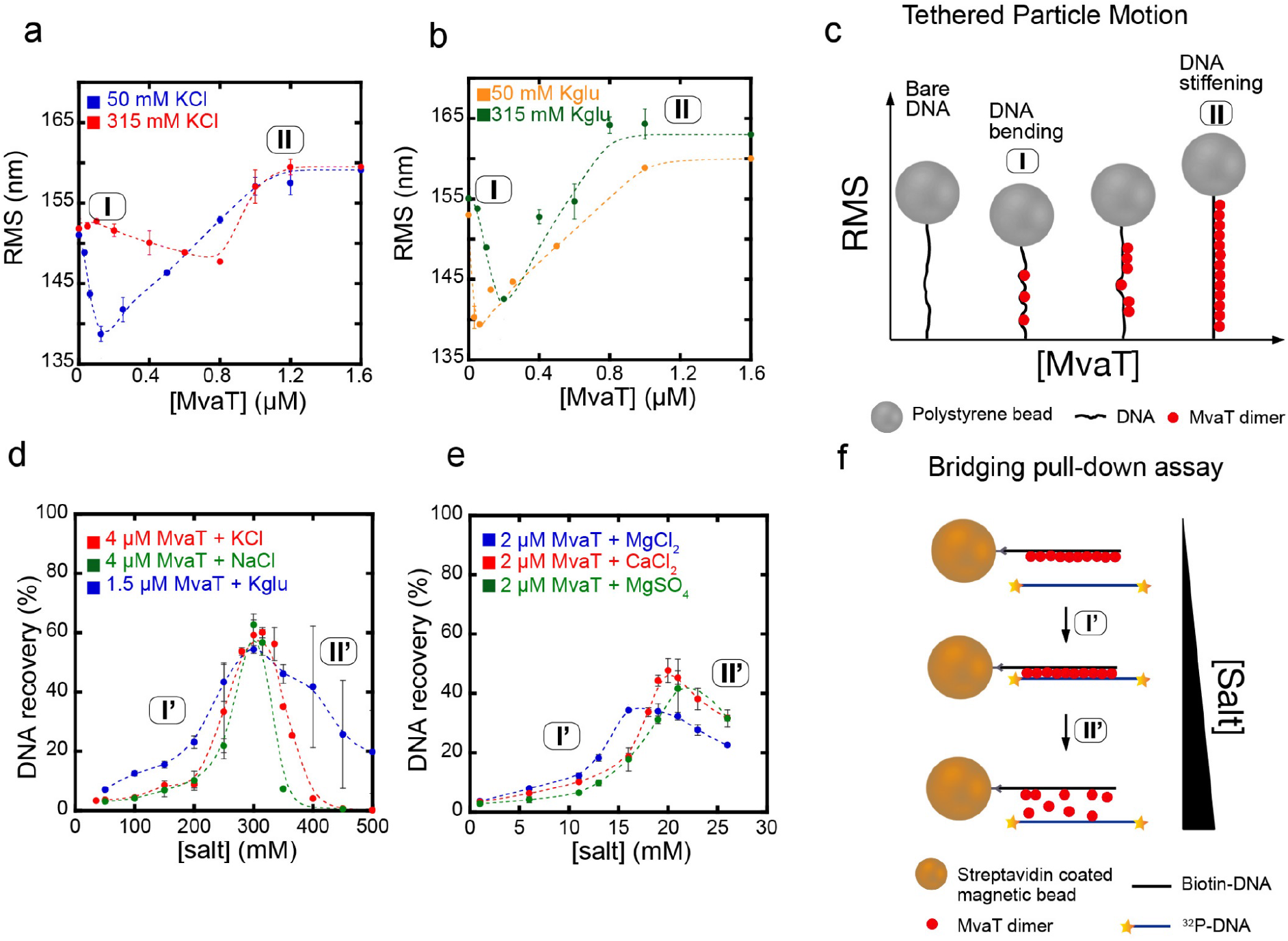
Conformational changes of MvaT induced by salt drive the switch between its DNA stiffening and bridging activities. (a) Root Mean Square displacement (RMS) of the tethered DNA particles as a function of MvaT concentration at 50 mM KCl (in blue) and 315 mM KCl (in red). The different regimes are labelled (I, II). (b) RMS as a function of MvaT concentration in the presence of 50 mM potassium glutamate (KGlu) (in yellow) and 315 m KGlu (in green). (c) Schematic representation of the experimental observations using the Tethered Particle Motion technique. The different regimes observed in (a) and (b) are labelled I and II. (d) DNA bridging efficiency of the MvaT wild type as a function of monovalent ions concentration: KCl (in red); NaCl (in green); KGlu (in blue). (e) The MvaT DNA bridging efficiency as a function of divalent ion concentration: MgCl_2_ (in blue); CaCl_2_ (in red) and MgSO_4_ (in green). The different regimes of MvaT DNA bridging activity are labelled (I’, II’). (f) Schematic representation of the experimental observations using the DNA bridging assay. The different regimes observed in (d) and (e) are labelled I’ and II’. Error bars indicate ± SD of duplicate.

Previously, the switch between DNA bridging and stiffening of H-NS family members was only observed and quantitatively analysed as a function of divalent ions ^19, 22^. Herein we showed that the DNA bridging efficiency of MvaT also depends on monovalent ions, suggesting that the switch might not require specific ions. There are some indications that H-NS can also bridge DNA in the presence of 200 mM KCl and in absence of MgCl_2_. While captured by AFM imaging, this observation was not corroborated by solution methods ^23^. To substantiate this observation, we here determined DNA bridging efficiency of H-NS in the presence of 50-200 mM KCl. Indeed, we observe a fourfold increase in the bridging efficiency between 50 and 100 mM KCl (fig. S5c). These findings indicate that the switch between the DNA binding modes of H-NS proteins is independent from the type of ions and is mainly due to the increase in ionic strength. Note, however that divalent ions are more efficient (by 1 order of magnitude) in promoting bridging than monovalent ions. These differences are possibly due to the higher effective shielding of the proteins charged regions by multivalent ions.

The observed effect of ionic strength on the ability of MvaT to bridge DNA can be explained structurally. The increase in salt concentration gradually destabilizes the interdomains electrostatic interactions of the MvaT protomers. DNA has an additive contribution on the shifting of the conformational equilibrium of the MvaT protomers towards the fully opened state. In this state, MvaT exposes most of the DNA binding domains in the lateral protein filament to concurrently interact with another DNA duplex and form a bridge. Beyond a certain threshold of ionic strength, the bridged protein-DNA complex starts disintegrating because the high ionic strength will weaken the interactions between the DNA binding domain of MvaT and DNA (fig. S5b).

These macroscopic observations provide direct evidence that the electrostatic interactions of the DBD and N-terminus of MvaT change, in response to altered ionic strength, translates into altered DNA binding properties. Salt and DNA function additively to disrupt these interdomains interactions, thus inducing the required conformational changes that drive the switch between the DNA bridging and stiffening activities of the H-NS family of proteins.

## Discussion

The mechanism by which H-NS family proteins sense the osmotic environment to modulate bacterial nucleoid organization and gene transcription is poorly understood. H-NS proteins form lateral nucleoprotein filaments along the DNA or bridge two DNA duplexes, resulting in DNA loops. Divalent ions drive the switch between DNA stiffening and bridging activities of the H-NS family members ^16, 19, 22^.

Our previous studies suggested that the switch between the two DNA binding modes requires H-NS protomers to experience conformational changes between an open and a closed state ^19^. Although the study was based on MD simulation and supported by mutagenesis, a detailed structural description of the protein conformational landscape based on experimental evidence was missing. To test this model, we have used MvaT from *P. aeruginosa*, which we argue represents a short structural version of H-NS as a benchmark.

Analysis of the primary sequences of H-NS family members has revealed conserved positions of charged residues despite the low sequence identity. A cluster of negatively charged residues is in the N-terminal domain whereas the DNA binding domain and the linker region contain positively charged residues. Thus, we hypothesised that electrostatic interactions between the two oppositely charges domains may take place, which could be modulated by the increase in ionic strength.

By combining PREs and SAXS techniques we have modelled the conformational landscape of the MvaT dimer under low and high salt conditions. Under both conditions, the protein samples heterogeneous conformational space where the two DNA binding domains adopt different spatial configurations. Two major conformers could be identified: a half-opened and a fully opened state. In the half-opened state, one of the DNA binding domains is interacting with the α2 N-terminal helix in an encounter complex like states. The interface between the two domains comprises the positively charged loop 95-ET**K**GGN**HK**-102 and the negatively charged segment 26-Q**DD**KLKK**E**L**EDEE**-38 of the DBD and the N-terminal α2, respectively. At low salt, these interactions prevent one of the DBDs of the MvaT protomer from forming a complex with DNA. At high salt, this electrostatic intramolecular complex is destabilized, causing an increase in exchange rate between the open and close states and thus a delicate change in the conformational space of the protein where it becomes more flexible and extended. This effect is enhanced by DNA substrate which induces the opening of the MvaT dimer by shielding the DBD from interacting with the N-terminal domain.

These findings agree with what has been proposed for H-NS. The MD simulations of the H-NS dimer in the presence of 50 mM KCl revealed interactions between its DBD and its N-terminal domain involving a negative amino acid patch in the α-helix 3 (E42-E50) and the positively charged regions 98–105 (KYSYVDENGE) and 123–129 (EQGKS) of the DBD ^19^. These interactions are altered in the presence of 130 mM KCl or 10 mM MgCl_2_. Recently, Arold and co-workers confirmed these findings by NMR titration of the N-terminal domain (2-57) with the DNA binding domain (84-137). The interdomain interactions were lost at high salt concentration. However, the authors claim that the occurrence of these interactions within the protein nucleofilament is not possible due to stereochemical restraints (as the linker is 25-30 Å from site 1) unless the oligomerization site 2 unfolds ^24^. We disagree with these claims as the linker region and the DNA binding domain together have a length of more than 40 Å, which provides the DBD with enough space and degree of freedom to reach the negative residues in α-helix 3. Additionally, disruption of the H-NS interdomain interactions by mutagenesis has generated a variant that adopts an open conformational state and oligomerizes, and is able to stiffen and bridge DNA in the absence of magnesium ^19^. This indicates that these interactions indeed occur in the H-NS oligomer and their modulation by salt or mutagenesis induces a switch between its DNA binding modes.

To establish structure-function relationships for the MvaT protomer we investigated the effects of salt on the DNA binding properties of MvaT using TPM and a pull-down bridging assay.

MvaT, like H-NS, oligomerizes along the DNA substrate, leading to stiffening of DNA, in agreement with previous reports ^37, 38^. Unlike H-NS, MvaT binds to the DNA substrate and induces bending. This is due to differences between the DNA binding mechanisms of the two proteins. MvaT binds to the DNA through an ‘AT-pincer’ motif which causes backbone bending ^40^. On the other hand H-NS binding to DNA involves an ‘AT-hook like’ structural motif ^14, 50^ that intercalates within the DNA helix without perturbing its twist ^51^.

H-NS bridges are disrupted at lower KCl concentration compared to MvaT, indicating a higher sensitivity to the ionic strength of the surrounding medium. These differences are likely due to differences in bridge stability resulting from the difference in DNA binding mechanisms, with MvaT forming a stronger bridge than H-NS. Additionally, H-NS requires a lower amount of MgCl_2_ compared to MvaT to reach the maximum bridging activity ^19^. This could be explained by stronger electrostatic interdomain interactions within MvaT compared to H-NS, thus requiring higher KCl and MgCl_2_ concentrations for their destabilization.

In contrast, StpA, an H-NS homologue, requires no or sub mM amounts of MgCl_2_ to be able to bridge DNA ^52, 53^. The high sensitivity of StpA towards magnesium chloride is possibly due to the weaker electrostatic interdomain interactions compared to H-NS and MvaT, as concluded from the charge distribution within its primary sequence (fig. S1).

These three examples indicate that although the H-NS family members might structurally respond in a similar way to the increase in ionic strength to switch between their DNA stiffening and bridging activities, variations on this theme exist. These differences are due to the delicate divergence in their intrinsic charge distribution, and thus the fine-tuning of the strength of their electrostatic interdomain interactions.

The recently proposed mechanism by which H-NS senses temperature to alleviate gene repression involves unfolding of site 2, which induces an autoinhibitory conformational change within the H-NS protomer^24^. Here we propose that H-NS family proteins do not require site 2 unfolding to sense the osmotic surrounding. Their osmosensitivity is mainly mediated by the modulation of the electrostatic interactions between their N-terminal and DNA binding domains within their nucleoprotein filaments. Altogether, we demonstrate that H-NS family proteins nucleofilaments might respond differently to the environmental changes in temperature and ionic strength.

Thus, based on the description of the conformational landscape of MvaT, we propose a general mechanism for the basic response of H-NS family proteins to changes in osmotic environment (fig. 6). Under low ionic strength, the H-NS protomers interact individually with AT-rich nucleation sites on the DNA. From these sites, the protein spreads cooperatively along the DNA substrate forming lateral nucleoprotein filaments where only one of the DNA binding domains of each dimer is engaged with the DNA substrate. The other domain is sequestered by electrostatic interactions with the N-terminal domain. The DNA substrate and the increase in ionic strengths additively destabilize these interactions. Consequently, the protomers release their second DNA binding domains to interact with another DNA duplex and form a bridge. Above a certain threshold of ionic strength, the bridge is dissociated due to loss of the electrostatic interactions that mediate the complex. In addition, changes in DNA topology induced by salt ^54^ might contribute to formation or rupture of bridges.

**Figure 6:**
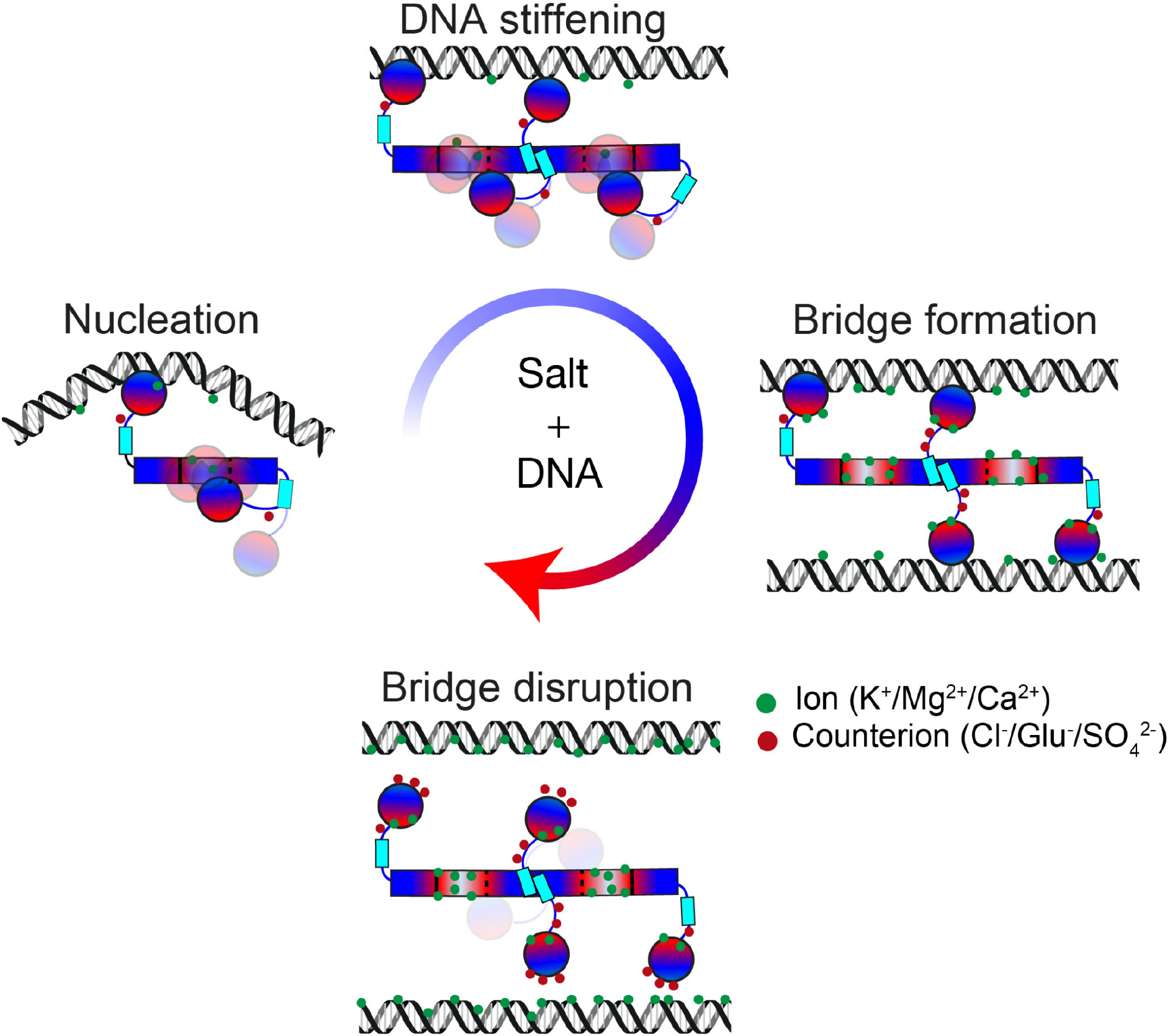
General mechanism of osmoregulation of the DNA binding properties of H-NS family proteins. The white, red and blue colours refer to neutral, negative and positive charged amino acid patch on the H-NS molecular surface.

In the present work, we have decrypted the molecular basis that governs DNA-DNA bridge formation by MvaT and proposed a structural dynamics paradigm for the osmosensitivity of H-NS family proteins. *In vivo* this mechanism is part of more extensive and complex regulatory network. Clearly this process is modulated by other regulatory factors, such as protein modulators of H-NS function, ions and their cellular concentrations, in conjunction with local DNA topology. In addition to a fundamental understanding of the activities of H-NS family proteins, our findings pave the way for the modulation of their DNA binding modes by small chemical compounds as a new generation of antibiotics.

## Material and Methods

### Construction of expression vectors and mutagenesis

The MvaT coding gene from P. *aeruginosa* and its DBD (residues 76-124) were subcloned into pET30b vector without and with C-terminal 6×His-tag, respectively, using Gibson assembly^55^. The Gibson assembly primers were purchased from Sigma-Aldrich. Mutated MvaT derivatives were generated using the same approach. Accuracy of the cloned sequences was confirmed by DNA sequencing.

### DNA substrates

All tethered particle motion and pull-down bridging assay experiments were performed using a random, AT-rich, 685 bp (32% GC) DNA substrate. The DNA substrate was generated by PCR and the products were purified using a GenElute PCR Clean-up kit (Sigma-Aldrich). If required, DNA was ^32^P-labeled as described previously ^19^. For the electrophoretic mobility shift assay an AT-rich 200 bp (32%GC) was generated using the same procedure.

The oligonucleotides to generate the 12 bp DNA duplex d(CGCATATATGCG) were purchased from Sigma Aldrich. The lyophilized oligonucleotides were solubilized in 20 mM Bis-Tris Buffer, 50 mM or 300mM KCl, pH 6, combined and hybridized by heating to 98°C for 3 min, followed by slow cooling down to room temperature. The 12 bp DNA duplex was used for NMR titration of the MvaT dimer and DBD.

### Protein expression and purification

*E. coli* BL21 competent cells were transformed with the MvaT pET30b vector and cultured in LB medium. For ^15^N ^13^C or ^15^N isotopic labelling, the transformed cells were first cultured in 2 mL LB medium and then transferred into 50 mL of inoculation culture in M9 medium (comprising the required isotopes) for incubation overnight at 37 °C. The inoculation culture was diluted in 1L M9 medium and was incubated at 37 °C until it reached an OD_600_ ≈ 0.6. MvaT production was induced by adding 100 μM of isopropyl β-D-1-thiogalactopyranoside (IPTG) and the culture was incubated overnight at 20 °C. The next day cells were harvested by centrifugation and the pellet was resuspended in the lysis buffer (20 mM Tris, 150 mM NaCl, 10 mM EDTA, pH 7). The cells were lysed using pre-cooled French press, and the lysate was centrifuged for 20 min at 8000 rpm. Next, the supernatant was loaded on a Heparin column (from GE healthcare Life sciences) and the protein was eluted by applying a NaCl gradient from 0.15 to 1 M. The eluted fractions were checked by SDS-PAGE and the ones that contain the MvaT dimer were pooled, concentrated and buffer changed using a PD10 column to 20 mM Bis-tris, 50 mM KCl, pH 6. Next, the protein was loaded on a SP column (from GE healthcare Life sciences) and eluted by applying a NaCl gradient (from 0.15 to 1 M). The eluted fractions were also checked by SDS-PAGE and the ones that contain the MvaT dimer were pooled and concentrated to a 500 μL volume with an Amicon 10 kDa cut-off filter. The concentrated protein fractions were loaded on a GE Superdex 75 10/300 GL column and eluted with 20 mM Bis-tris, 150 mM KCl, pH 6. The collected fractions were buffer exchanged to either 20 mM Bis-tris, 50 mM KCl or 300mM KCl, 1mM EDTA, pH 6. The purity of the protein was checked by SDS-PAGE (>95%) and the concentration was determined using a bicinchoninic acid assay.

The same procedure was used for the DBD over-expression and purification. The only difference in purification was in the first step where a nickel column was used instead of a heparin column.

### The Tethered Particle Motion (TPM)

The Tethered Particle Motion experiments were done as previously described ^19, 56^ and the flow cells were prepared with minor modifications. In short, first, the flow cell was washed with 100 μl of wash buffer (10 mM Tris-HCl pH 8, 150 mM NaCl, 1 mM EDTA, 1 mM DTT, 3% Glycerol, 100 μg/ml BSA (ac)) and experimental buffer (10 mM Tris pH 8, 10 mM EDTA, 5% glycerol with different salt concentrations as indicated in the text). The flow cell was sealed after adding protein (0 nM to 1600 nM) in experimental buffer, followed by 15 minutes of incubation. The measurements were initiated 15 min after introducing protein. For each flow cell more than 200 beads were measured at a temperature of 25°C. Data analysis was done as reported previously ^19, 56^.

### Pull down bridging Assay

The DNA Bridging Assay was performed as described previously ^19, 49^, with minor modifications. The Streptavidin-coated Magnetic Dynabeads M280 (3 μL per sample) were resuspended in 6 μl coupling buffer (CB: 20 mM Tris-HCl pH 8.0, 2 mM EDTA, 2 M NaCl, 2 mg/mL BSA (ac), 0.04% Tween20) containing 3 μl 100 fmol biotinylated 32% GC DNA (685bp), and the bead suspensions were incubated for 20 min at 25°C on a shaker (1000 rpm). The beads were then washed twice with incubation buffer (IB: 10 mM Tris-HCl pH 8.0, 5% v/v Glycerol, 1 mM Spermidine) before resuspension in IB. Next, salt (KCl, NaCl, Kglu, CaCl_2_, MgCl_2_ or MgSO_4_) was added to the desired concentration in each sample. Radioactive DNA (around 8000 cpm) was added to each sample and unlabeled 685 bp (32% GC) DNA was added to maintain a constant (2 fmol/μl) DNA concentration. MvaT or H-NS proteins were added to initiate the formation of bridged DNA-DNA complex. The mixture was incubated for 20 min (unless indicated otherwise) on the shaker at 1000 rpm at 25°C. After the incubation, the beads of each sample were washed once with its experimental buffer, before resuspension in 12 μl counting buffer (10 mM Tris pH 8.0, 1 mM EDTA, 200 mM NaCl, 0.2% SDS). The radioactive signal of samples was quantified by liquid scintillation over 5 min. Sample values recovered from the bridging assay were corrected for background signal (from a control sample without protein). The DNA recovery was calculated based on a reference sample containing the amount of labeled ^32^P 685 bp DNA used in each sample and expressed as ‘DNA recovery (%)’. All DNA bridging experiments were performed at least in duplicate. Each experiment contained 20 mM KCl, to which additional salt (KCl, NaCl, Kglu, CaCl_2_, MgCl_2_ or MgSO_4_) was added depending on the tested experimental conditions.

### Electrophoretic mobility shift assay

The assay was performed on an AT-rich 200 bp DNA substrate (32%GC). Typically, 25 ng of DNA was incubated with different concentrations of the MvaT wild type or the MvaT dimer F36D/M44D in the binding buffer (10 mM Tris-HCl, 50 mM KCl, pH 7.5). The mixture was loaded on a 1% agarose gel (containing 1:10^4^ dilution of Gel red DNA stain) with 1μl of 6 × loading buffer (10mM Tris-HCl pH 7.6, 0.03% bromophenol blue, 0.03 % xylene cyanol FF, 60% glycerol and 60 mM EDTA). The samples were run at 16 mA for 4 hours at 4°C in TAE standard buffer. The gels were visualized with a Bio-rad Gel Doc XR+ system.

### Circular dichroism

The MvaT wild type and F36D/M44D dimer far-UV CD spectra were recorded on a Jasco 810 CD spectrometer using a 10 μM protein solution in 20 mM Bis-Tris Buffer, 150 mM KCl, pH 6. The spectra were recorded in a 0.1 cm path length quartz cuvette using four scans with a bandwidth and a wavelength step of 1 nm. The obtained spectra were background corrected and smoothed using Jasco Spectra Manager.

### Isothermal titration calorimetry

ITC experiments were performed using a MicroCal VP-ITC system at 20 °C. The samples contained 20 mM Bis-Tris, pH 6 with 50 mM KCl. Typically, 10 μM of MvaT F36D/M44D was placed in the cell (1.4 mL) and titrated with 360 μM (500 μL volume) of the DNA duplex d(CGCATATATGCG), injected in 2 μL aliquots. The delay time between the injections was 60 s with a stirring speed of 307 rpm. The corresponding “protein to buffer” controls were performed for background correction. The ITC titration data were analysed using Origin 7.0 (OriginLab) provided with the instrument. Standard deviation was calculated from the fit by Origin.

### Small angle X-ray scattering

All the experiments were performed at the ESRF BioSAXS beamline BM29, Grenoble, France ^57^. Given the sensitivity of the batch SAXS mode for even small amounts of large soluble aggregates we used SEC-SAXS approaches to measure SAXS data only on monodisperse samples. A volume of 250 μL of protein for each MvaT sample was loaded on a GE Superdex 75 10/300 GL column *via* a high performance liquid chromatography (HPLC) system (DGU-20A5R, Shimadzu, France) attached directly to the sample-inlet valve of the BM29 sample changer. Low salt MvaT sample was measured at 11 mg/mL in 20 mM Bis-Tris, 50 mM KCl, pH 6.0, while high salt MvaT sample was measured at 9.5 mg/ mL in 20 mM Bis-Tris, 300 mM KCl, pH 6.0. The column was equilibrated with 3 column volumes to obtain a stable background signal, which was confirmed before measurement. All the SAXS data were collected at a wavelength of 0.99 Å using a sample-to-detector (PILATUS 1 M, DECTRIS) distance of 2.81 m. The scattering of pure water was used to calibrate the intensity to absolute units ^58^. Data reduction was performed automatically using the EDNA pipeline ^59^. Frames in regions of stable *R*_*g*_ were selected and averaged using PRIMUS ^60^ to yield a single averaged frame corresponding to the scattering of individual SEC species. Analysis of the overall parameters was carried out by PRIMUS from ATSAS 2.8.4 package ^61^ and by ScÅtter 3.0 software. The pair distance distribution function, P(r), and maximum diameter of the particle (*D*_*max*_) were calculated in GNOM using indirect Fourier transform method ^62^. The information on data collection and derived structural parameters is summarized in Table S1. The pair distance distribution functions were used to calculate *ab initio* models in P1 symmetry with DAMMIN; the models were averaged, aligned and compared using DAMAVER ^61^.

### NMR resonance assignments

Sequential assignment of the MvaT dimer at 20 °C was performed using the triple resonances HNCACB, CBCAcoNH, HNCA, HNcoCA, HNCO and HNcaCO experiments on a ^15^N, ^13^C MvaT NMR sample (0.5 mM in 20 mM Bis-Tris, 150 mM KCl, 1mM EDTA, pH 6). The NMR spectra were recorded on a Bruker Avance III (HD) 600 MHz spectrometer, equipped with TCI cryoprobe, processed by TopSpin 3.5 (Bruker Biospin) and analysed with Sparky software^63^. The peak list of MvaT dimer was used to assign the MvaT dimer K31C mutant HSQC spectrum.

### NMR titration experiments

The NMR titration of the MvaT dimer and DBD with the 12 bp DNA substrate was performed on a 95 or 100 μM ^15^N isotopically labelled protein samples, respectively, in 20 mM Bis-Tris Buffer, 50 mM KCl or 300 mM KCl, pH 6 and 6% D_2_O. A series of ^1^H-^15^N HSQC spectra were acquired by gradually increasing the DNA:MvaT molar ratio from 0.1 or 0.2 to 1.6. The experiments were recorded at 20 °C on a Bruker Avance III (HD) 600 MHz spectrometer, equipped with TCI cryoprobe, processed by TopSpin 3.5 (Bruker Biospin) and analysed by CCPnmr software^64^. The MvaT dimer titration with KCl was conducted using the same procedure. The reference titration points were collected in a solution with 20 mM Bis-Tris, 50 mM KCl, pH 6 and the potassium chloride concentration was increased in steps to 100, 150, 200, 250 and 300 mM KCl.

The changes in peak positions and intensities were analysed by an in-house python script and the average chemical shift perturbations (CSP) were calculated using equation (1):

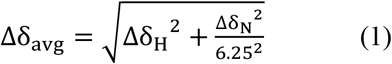

Line shape analysis of the titration data was performed by TITAN ^65^ using a two-state binding model.

### Protein labelling and PRE measurement

To attach the MTS [(1-Acetyl-2,2,5,5-tetramethyl-3-pyrroline-3-methyl)-methanethiosulfonate] and MTSL [S-(1-oxyl-2,2,5,5-tetramethyl-2,5-dihydro-1H-pyrrol-3-yl)methyl methanesulfonothioate] spin labels (bio-connect Life sciences) on the MvaT dimer surface, K31 was substituted with a cysteine. A ^15^N isotopically labelled sample of the variant MvaT K31C/F36D/K44D was produced using the procedure described above. The purified protein was first incubated for 1 hour at room temperature in the presence of 10 mM DTT to reduce disulphide bonds, followed by buffer exchange using PD10 desalting column. The protein was directly eluted into a solution of 20 times molar excess of MTS or MTSL. The mixture was incubated for 1 h stirring at 4 °C. The sample was concentrated to a volume of 500 μL with an Amicon 10 kDa cut-off filter and injected into a GE Superdex 75 10/300 GL column to remove excess spin label. Incorporation of MTSL was >95% according to mass spectrometry, and the dimeric state of the conjugated protein was checked by analytical size exclusion chromatography. PREs were obtained using ^1^H-^15^N TROSY spectra of 185 μM MvaT in 20 mM Tris, pH 6 at low salt (50 mM KCl) or high salt (300 mM) and in the presence or absence of DNA, with a DNA-to-protein molar ratio of 1.6. The TROSY spectra of the diamagnetic and paramagnetic samples were processed using TopSpin 3.5 and peak intensities and line widths were extracted using CCPnmr analysis^64^. The paramagnetic contribution to the transverse relaxation rate (R2sp) was estimated from the measured intensity ratio of the paramagnetic and diamagnetic samples using equation^66^ (2):

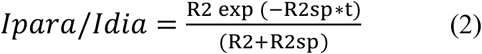

where I_para_/I_dia_ is the measured intensity ratio, R_2_ is the relaxation rate estimated from the widths at the half-height of the diamagnetic sample resonances, and t is the time during which ^1^H magnetisation is in the transverse plane during the TROSY pulse sequence (10 ms).

### MvaT structure homology modelling

The full-length MvaT dimer was constructed by combining a homology model of the dimerization domain (residues 1-78) and the NMR structure of the DNA binding domain (PDB ID: 2MXE) ^40^ (residues 79-124), using Modeller version 9.15. The homology model for the dimerization domain was obtained by aligning residues 1-78 of MvaT to the crystal structure of the N-terminal domain of *P. putida* TurB (PDB ID: 5B52) ^39^.

### Conjoint ensemble refinement against PRE and SAXS data

All simulations were performed in Xplor-NIH^67, 68^, starting from the homology model of the MvaT F36D/M44D dimer generated in this work (see text). The positions of the N-terminal domains of the MvaT dimer (residues 1-47) were fixed, the C-terminal DNA-binding domain (residues 83-122) was treated as a rigid-body group, while the intervening linker was given full degree of freedom. The computational protocol comprised an initial rigid-body simulated annealing step followed by the side-chain energy minimization as described before^69^. The energy function consisted of standard geometric (bonds, angles, dihedrals, and impropers) and steric (van der Waals) terms, a knowledge-based dihedral angle potential, and the PRE and SAXS energy terms incorporating the experimental data^70^. Truncated SAXS curves (q < 0.4 Å^−1^) and the PRE data for the C-terminal DNA-binding domain and the interdomain linker (residues 50-124) at each salt condition were used as the experimental input^69, 70^. Multiple copies of MvaT dimers (N = 1-5) were refined simultaneously in order to simulate molecular ensembles of multiple conformers^69^ (Note that this procedure allows for the atomic overlap among MvaT molecules constituting an ensemble). In each run, 100 independent calculations were performed, and 10 lowest-energy solutions were selected for further analysis.

During the simulations, the PRE contributions from both subunits were taken into account. The PRE for each residue was defined as the sum of four contributions: 1) PRE from spin label (SL) on subunit 1 to the residue on subunit 1; 2) PRE from SL on subunit 1 to the residue on subunit 2; 3) PRE from SL on subunit 2 to the residue on subunit 2; 4) PRE from SL on subunit 2 to the residue on subunit 1.

To assess the agreement between the observed PREs and the PREs back-calculated from the simulated ensembles generated in each run, we calculated the *Q* factor^71^, equation (3):

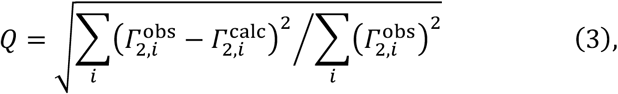

where 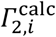 is given by equation (4):

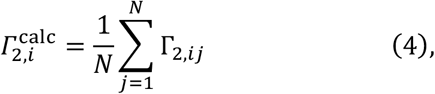

where *N* is the ensemble size and Γ_2,*i,j*_ is the PRE back-calculated for the amide proton of residue (*i*) of the ensemble member (*j*).

The agreement with the experimental SAXS data in each run was assessed by calculating χ^2^ with the calcSAXS helper program^69^, which is part of the Xplor-NIH v 2.49 package used in this work, assuming equal conformer populations in the simulated ensembles. The lowest energy structures of the MvaT dimer obtained from the ensemble modelling were fitted into the *ab initio* molecular shapes obtained by DAMMIN using UCSF Chimera software ^72^.

## Acknowledgments

This research was supported by grants from the Netherlands Organization for Scientific Research [VICI 016.160.613] (R.T.D.), the Human Frontier Science Program (HFSP) [RGP0014/2014] (R.T.D.) and a grant from the China Scholarship Council (CSC) [201506880001] (L.Q.). We thank Dr. Navraj Pannu for establishing the contact with Dr. Gabriele Giachin at the European Synchrotron Radiation Facility to perform the SAXS experiment. Maud Bremer is acknowledged for making structural models.

## Contributions

L.Q., F.B.B., G.G. and P.V.S. performed the experiments. L.Q., F.B.B., Y.G.J.S., A.N.V., J.V., G.G., M.U. and R.T.D. contributed to data analysis and discussion. F.B.B., M.U. and R.T.D. supervised the project. F.B.B. and R.T.D. wrote the manuscript. All authors reviewed and corrected the manuscript.

