## Supplemental Material for "Structural basis for osmotic regulation of the DNA binding properties of H-NS proteins"

[1] Department of Macromolecular Biochemistry, Leiden Institute of Chemistry, Einsteinweg 55, 2333 CC Leiden, the Netherlands
[2] Centre for Microbial Cell Biology, Leiden University, Einsteinweg 55, 2333CC Leiden, The Netherlands
[3] Laboratory of Medical Biochemistry, University of Antwerp, Campus Drie Eiken, University Square 1, 2610 Wilrijk, Belgium
[4] VIB-VUB Structural Biology Research Center, Pleinlaan 2, 1050 Brussels, Belgium
[5] Jean Jeener NMR Centre, VUB, Pleinlaan 2, 1050 Brussels, Belgium
[6] Department of Computational Chemistry, Van't Hoff Institute for Molecular Sciences, University of Amsterdam Science Park 904, 1098 XH, Amsterdam, The Netherlands
[7] Structural Biology Group, European Synchrotron Radiation Facility (ESRF), Grenoble, France.
[◊]Contributed equally to this work

[⊥]Corresponding authors:
Fredj Ben Bdira:
Remus T. Dame:

**Keywords:**
H-NS family proteins; MvaT; interdomains electrostatic interactions; osmosensitivity; DNA bridging.

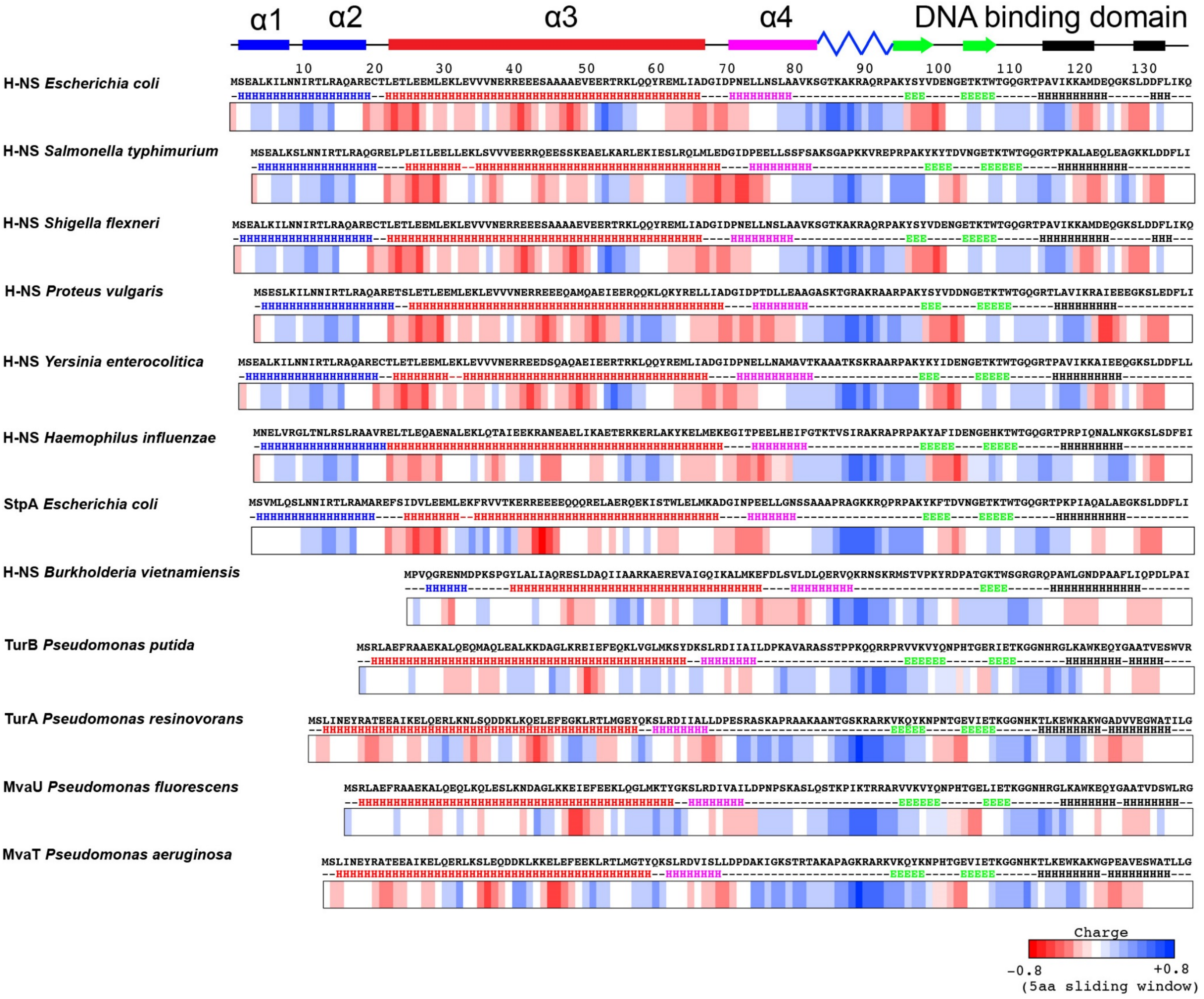

**Fig. S1:** Fold topology and charged residue/charge distributions of H-NS family proteins. Sequence alignment of twelve H-NS family proteins from different bacterial organisms was generated using Clustal omega[1]. The multi-alignment file was used to predict the secondary structures content using JPRED server [2]. (H) is for α-helices, (B) for β-sheets, (-) for random coils. A schematic of the *E. coli* H-NS secondary structure is depicted above. The lower panels represent the 5 amino acids average charge of the primary sequences of H-NS family members. Positively and negatively charged regions are colored with blue and red rectangles, respectively. The neutral amino acid patches are in white.

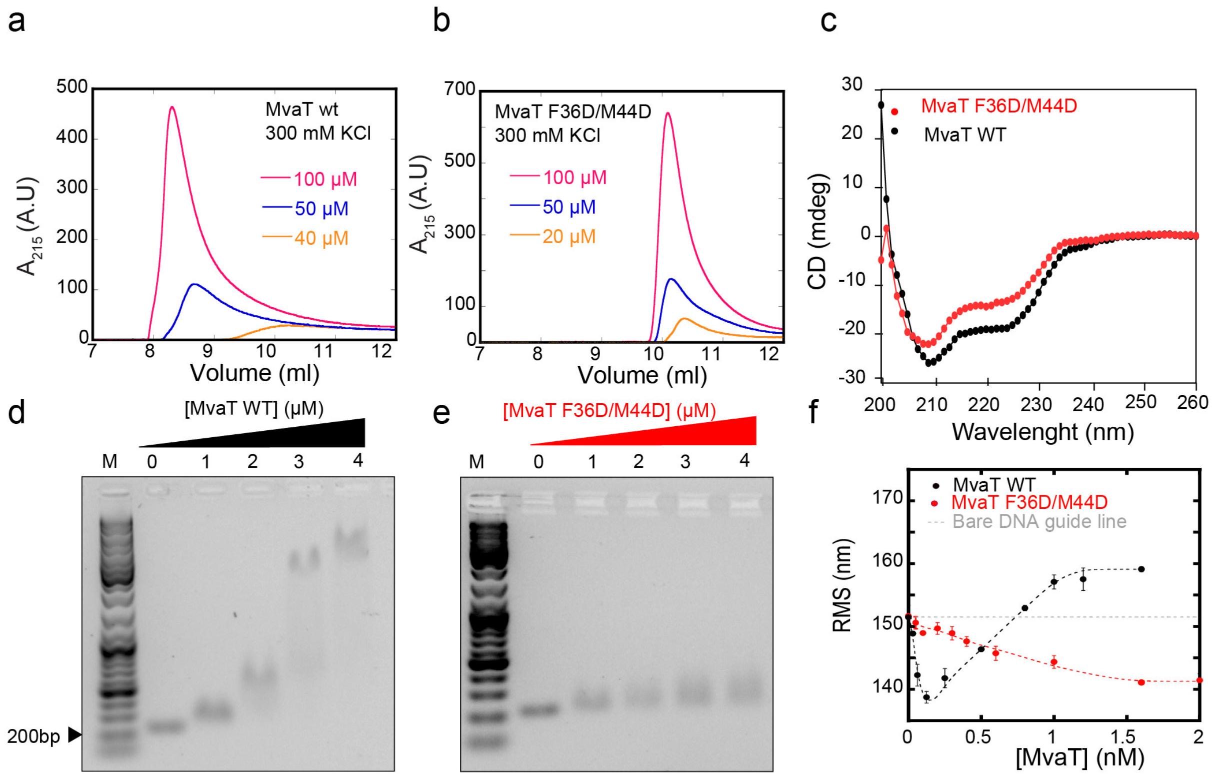

**Fig. S2:** The effects of F36D/M44D double mutations on MvaT secondary/quaternary structure and DNA binding. (a) and (b) concentration dependency of the MvaT wild type and the F36D/M44D mutant oligomerisation state in the presence of 300 mM KCl as monitored by size exclusion chromatography. (c) Overlay between the far-UV CD spectra of MvaT WT (in black) and MvaT F36D/M44D (in red). (d) and (e) Electrophoretic mobility shift assay on a 1% agarose gel of MvaT WT and MvaT F36D/M44D using a 200 bp AT-rich DNA substrate (32% GC), respectively. (f) Overlay of RMS date obtained for titration of MvaT F36D/M44D (red line) and MvaT wild type (black line) at 50 mM KCl in TPM experiments. The dashed grey line indicates the RMS of bare DNA.

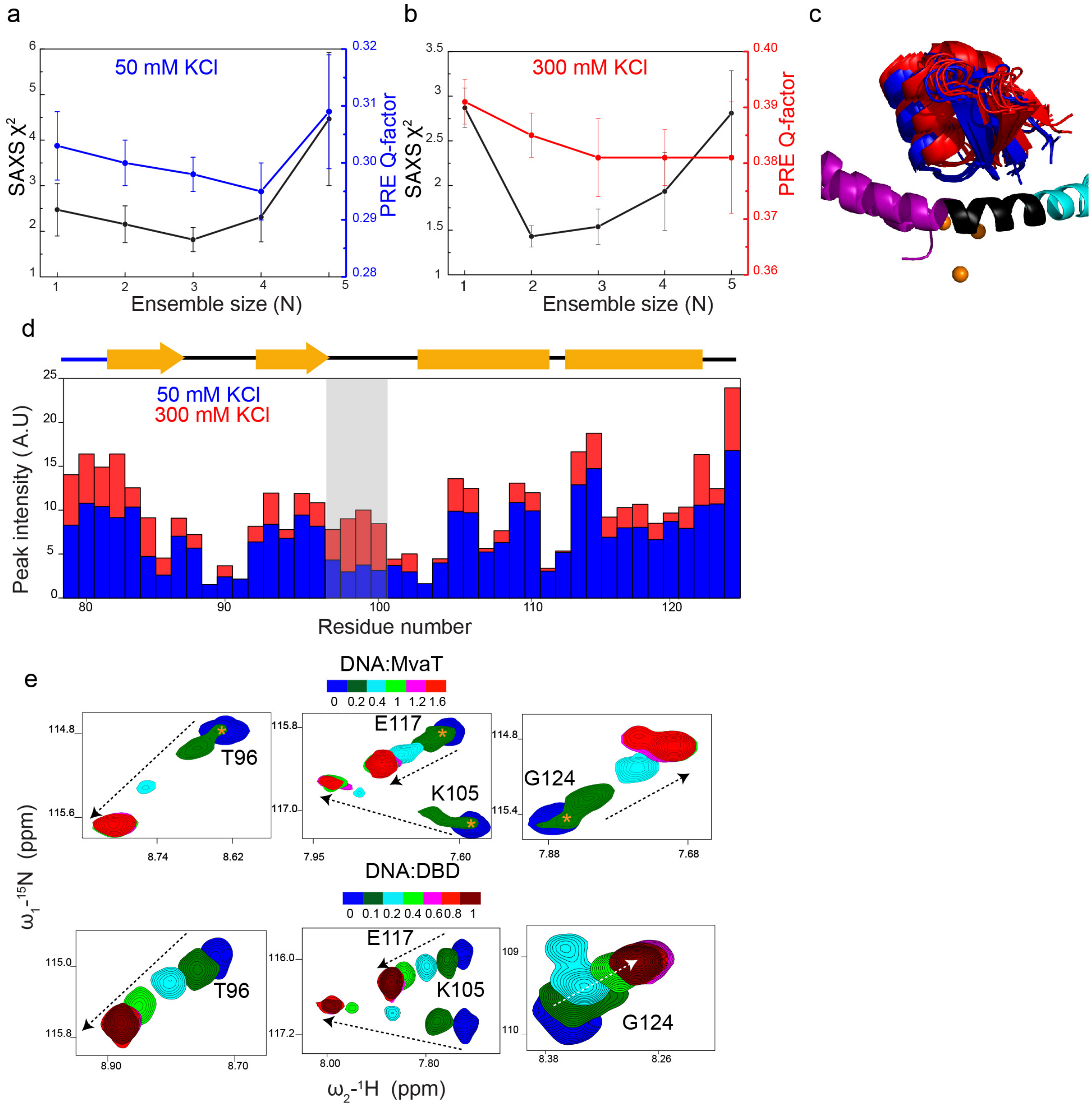

**Fig. S3:** MvaT structural ensemble calculation quality metrics and the DNA binding mechanism of the MvaT dimer and individual DBD. (a) and (b) show the variation of the SAXS $\chi^2$ and PREs Q-factor versus the ensemble size (N) of the low and high salt MvaT structural modelling, respectively. (c) Shows the position of the DNA binding domain relative to the N-terminal domain of the lowest energy structures at low (blue cartoon) and high salt (red cartoon). The orange spheres represent the nitroxide positions determined within the ensemble structure calculation. (d) MvaT dimer DBD peak intensities at 50 mM KCl (blue bars) and 300 mM KCl (red bars). Loop 95-102 is highlighted with the grey rectangle. (e) CSP amplitudes and directions (dashed arrows) of the MvaT dimer DBD resonances (upper panels) and the individual DBD with C-terminal 6×His-tag (lower panels) upon titration with 12 bp DNA performed in 20 mM Bis-Tris, 50 mM KCl, pH 6, molar ratios are indicated. The resonances which correspond to the N-terminal bound DBD are indicated by orange asterisks.

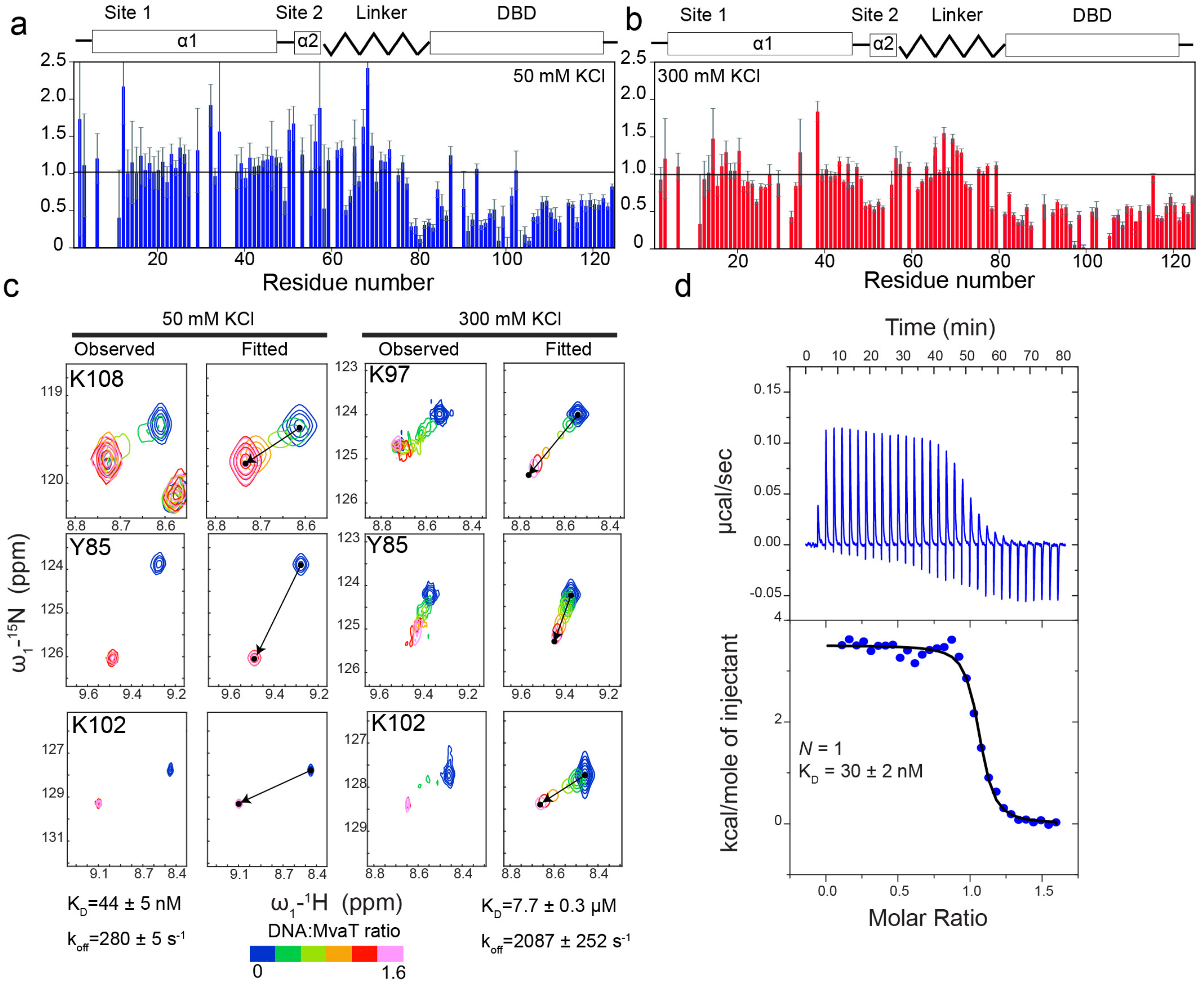

**Fig. S4:** MvaT F36D/F44D titration with 12 bp DNA AT rich DNA substrate. (a) and (b) changes in the MvaT dimer HSQC peak intensities ratio between at 1.6 DNA:MvaT molar ratio at 50 and 300 mM KCl salt concentration versus residue number, respectively. The layout of the MvaT secondary structure is shown in the upper panel. (c) MvaT dimer HSQCs line shape analysis upon titration with the 12 bp DNA substrate at low and high salt concentration. Experimental and fitted resonances of selected residues are shown. (d) ITC titration of the MvaT dimer with 12 bp DNA substrate at 50 mM. The extracted binding constant are indicated.

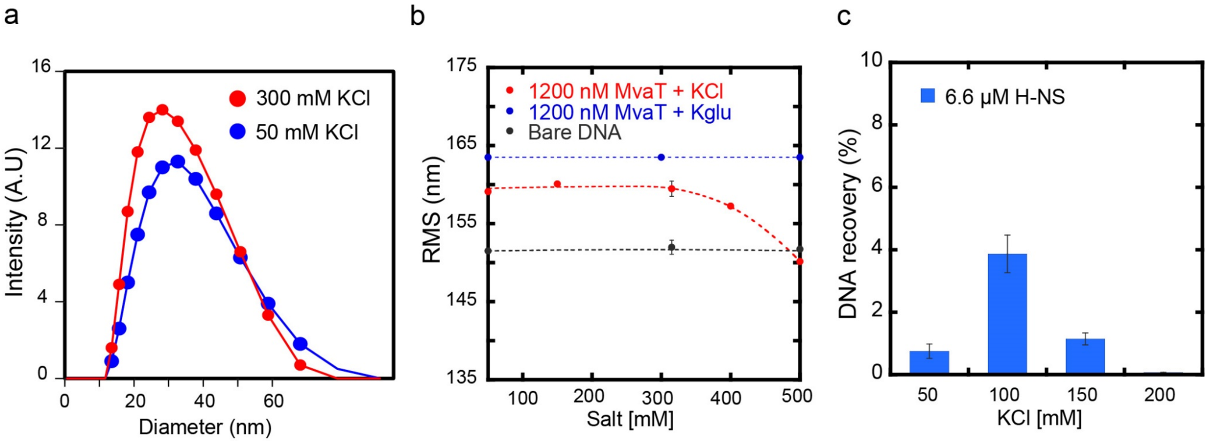

**Fig. S5:** Salt effect on H-NS protein oligomerization, stiffening and bridging activities. (a) MvaT wild type oligomer particle size at 100 μM concentration in the presence of 300 mM KCl (blue curve) and 50 mM KCl (red curve) determined by dynamic light scattering. (b) Effect of KCl (in red) and KGlu (in blue) concentration on MvaT wild type DNA stiffening activity using TPM. The effect of salt concentration on the RMS of the bare DNA is shown in black dashed line. (c) KCl concentration effect on H-NS DNA bridging efficiency using the DNA bridging assay. Error bars indicate the ± SD of duplicates.

**Table S1.** SAXS data collection and scattering-derived parameters.

| | **MvaT (low salt)** | **MvaT (high salt)** |
|---|---|---|
| **Data collection parameters** | | |
| Beam line | BM29 (ESRF) | BM29 (ESRF) |
| Wavelength (Å) | 0.99 | 0.99 |
| *q* range ($Å^{-1}$)* | 0.003 - 0.493 | 0.003 - 0.493 |
| Concentration (mg $ml^{-1}$) (mode) | 11 (SEC-SAXS) | 9.5 (SEC-SAXS) |
| Buffer conditions | 20 mM Bis-Tris, 50 mM KCl, pH 6.0 | 20 mM Bis-Tris, 300 mM KCl, pH 6.0 |
| Temperature (°C) | 20 | 20 |
| **Structural parameters**§ | | |
| *I(0)* ($cm^{-1}$) [from Guinier] | 69.81 ± 0.11 | 74.46 ± 0.01 |
| $R_g$ (Å) [from Guinier] | 35.63 ± 0.78 | 38.29 ± 0.65 |
| *I(0)* ($cm^{-1}$) [from *p(r)*] | 70.31 ± 0.10 | 74.89 ± 0.08 |
| $R_g$ (Å) [from *p(r)*] | 37.19 ± 0.11 | 39.89 ± 0.09 |
| $D_{max}$ (Å) | 146.99 | 158.38 |
| Porod volume estimate, $V_p$ ($Å^3$) | 47235 | 49908 |
| Porod exponent | 3.2 | 2.7 |
| **Molecular mass determination** | | |
| MM (kDa) [from *SAXSMoW* on final merged curve] | 31.50 ($q$ = 0.4 $Å^{-1}$) | 29.2 ($q$ = 0.3 $Å^{-1}$) |
| MM (kDa) [from $Q_R$ on final merged curve] | 28.09 ($q$ = 0.4 $Å^{-1}$) | 26.45 ($q$ = 0.3 $Å^{-1}$) |
| MM (kDa) [from $V_p$/1.7] | 27.78 | 29.35 |
| Calculated MM from sequence (kDa) | 28.26 | 28.26 |
| **Software employed** | | |
| Data processing and analysis | ATSAS, Scatter | ATSAS, Scatter |
| Computation of theoretical intensities and fitting | CRYSOL | CRYSOL |

Abbreviations: *I(0)*, extrapolated scattering intensity at zero angle; $R_g$, radius of gyration calculated using either Guinier approximation (from Guinier) or the indirect Fourier transform package GNOM [from *p(r)*]; *MM*, molecular mass; $D_{max}$, maximal particle dimension; $V_p$, Porod volume; $V_{ex}$, particle excluded volume

§Values reported are for average of frames in the peak for both MvaT samples.

*Momentum transfer $|q| = 4\pi\sin(\theta)/\lambda$

1. Sievers F*, et al.* Fast, scalable generation of high-quality protein multiple sequence alignments using Clustal Omega. *Mol. Syst. Biol.* **7**, 539 (2011).

2. Drozdetskiy A, Cole C, Procter J, Barton GJ. JPred4: a protein secondary structure prediction server. *Nucleic Acids Res.* **43**, W389-W394 (2015).

3. Waudby CA, Ramos A, Cabrita LD, Christodoulou J. Two-Dimensional NMR Lineshape Analysis. *Sci Rep.* **6**, 24826 (2016).
